# CD36 phosphorylation alters the thrombospondin binding site and reduces internal cavity accessibility and volume

**DOI:** 10.64898/2026.08.25.747030

**Authors:** Golshad Ghojoghi, Sylvain Chemtob, William D. Lubell, Huy Ong, Deniz Meneksedag-Erol

**Affiliations:** Department of Chemical and Materials Engineering, Gina Cody School of Engineering and Computer Science, Concordia University, Montreal, QC, Canada; Department of Chemistry & Biochemistry, Concordia University, Montreal, QC, Canada; Program in Molecular Biology, Faculty of Medicine, Université de Montréal, Montreal, QC, Canada; Department of Pediatrics, Sainte-Justine University Hospital Research Center, Université de Montréal, Montreal, QC, Canada; Department of Chemistry, Université de Montréal, Montreal, QC, Canada; Faculty of Pharmacy, Université de Montréal, Montreal, QC, Canada

**Author notes:** Corresponding author (DME).

## Abstract

The cluster of differentiation 36 (CD36) is a membrane protein with broad physiological roles in health and disease, and its function is regulated in part by phosphorylation. Experimental evidence shows that phosphorylation of Thr92 reduces CD36 affinity for thrombospondin-1 (TSP-1), binding of which initiates antiangiogenic signaling, whereas phosphorylation of Ser237 decreases CD36-mediated fatty acid uptake, with implications for energy metabolism. However, the only available crystal structure of CD36 lacks phosphorylation, and the molecular mechanisms by which phosphorylation regulates CD36 function remain largely unknown. This study provides an atomically detailed computational characterization of CD36 in unphosphorylated and dual phosphorylated states, using molecular dynamics simulations with a total sampling time of 30 µs in combination with Markov state models. We present, to our knowledge, the first evidence of a cryptic pocket on CD36 surface that is formed by phosphorylation. This cryptic surface pocket and a loop spanning residues 121-131 form a high affinity binding site for TSP-1 derived ligands, shifting their binding away from the canonical site. We propose that this altered binding provides a molecular basis for the disruption of antiangiogenic signaling upon CD36 phosphorylation. Additionally, our data indicate that, phosphorylation increases helicity and compaction within the helix-loop region spanning residues 296–331, narrowing one of the entrances to the internal cavity and reducing its overall volume. These conformational changes provide a potential mechanistic explanation for the decrease in fatty acid uptake upon CD36 phosphorylation. Our findings provide structural insights that may inform the future design of CD36 modulators and emphasize the importance of targeting phosphorylation induced CD36 conformations in angiogenic and metabolic diseases.

## Introduction

The cluster of differentiation 36 (CD36) is a multifunctional, membrane-associated protein with a central role in health and significant therapeutic potential across various diseases. CD36 is expressed in a wide range of cells and tissues including platelets, cells of the innate immune system, cardiovascular system, epithelial cells of the retina, breast and intestine, as well as adipose tissue and skeletal muscle [see reviews (Febbraio, Hajjar, and Silverstein 2001) and (Febbraio and Silverstein 2007)]. It binds a wide variety of ligands and regulates lipid metabolism (Abumrad et al. 1993), energy production (Manio et al. 2017), cellular adhesion (Leung et al. 1992), inflammation (Stewart et al. 2010), and immune response (Urban, Willcox, and Roberts 2001) in a cell and tissue specific manner. CD36 undergoes several posttranslational modifications, such as phosphorylation and glycosylation of its extracellular domain, which in part regulate its ligand binding and function. Disruption in CD36 signalling or expression is implicated in a number of diseases including age-related macular degeneration (Dorion et al. 2021), Alzheimer’s disease (El Khoury et al. 2003), cardiovascular illnesses (Ramos-Arellano et al. 2014), diabetes (Griffin et al. 2001), and cancer (Hale et al. 2014). Despite substantial evidence highlighting the therapeutic potential of CD36, its conformational dynamics remain poorly understood at the atomic level.

Current structural information on CD36 is limited to a single crystal structure of its glycosylated extracellular domain (PDB ID: 5LGD) (Hsieh et al. 2016), which lacks acetylation and phosphorylation. The extracellular domain of CD36 comprises 410 residues and contains the multiligand binding site (residues 139-183) (Glatz and Luiken 2018), binding sites for thrombospondin-1 (TSP-1) (CLESH: CD36 LIMP-II Emp sequence homology domain, residues 93-120) (Pearce, Wu, and Silverstein 1995) and oxidized low-density lipoprotein (oxLDL, residues 155-183) (Kar et al. 2008; Navazo et al. 1996) (Figure 1). Experimental evidence indicates that CD36 has two phosphorylation sites in its extracellular domain in residues threonine 92 (Thr92) and serine 237 (Ser237) (Figure 1b), and that phosphorylation at these sites affects ligand binding and fatty acid uptake, respectively (Chu and Silverstein 2012; Guthmann et al. 2002).

**Figure 1.**
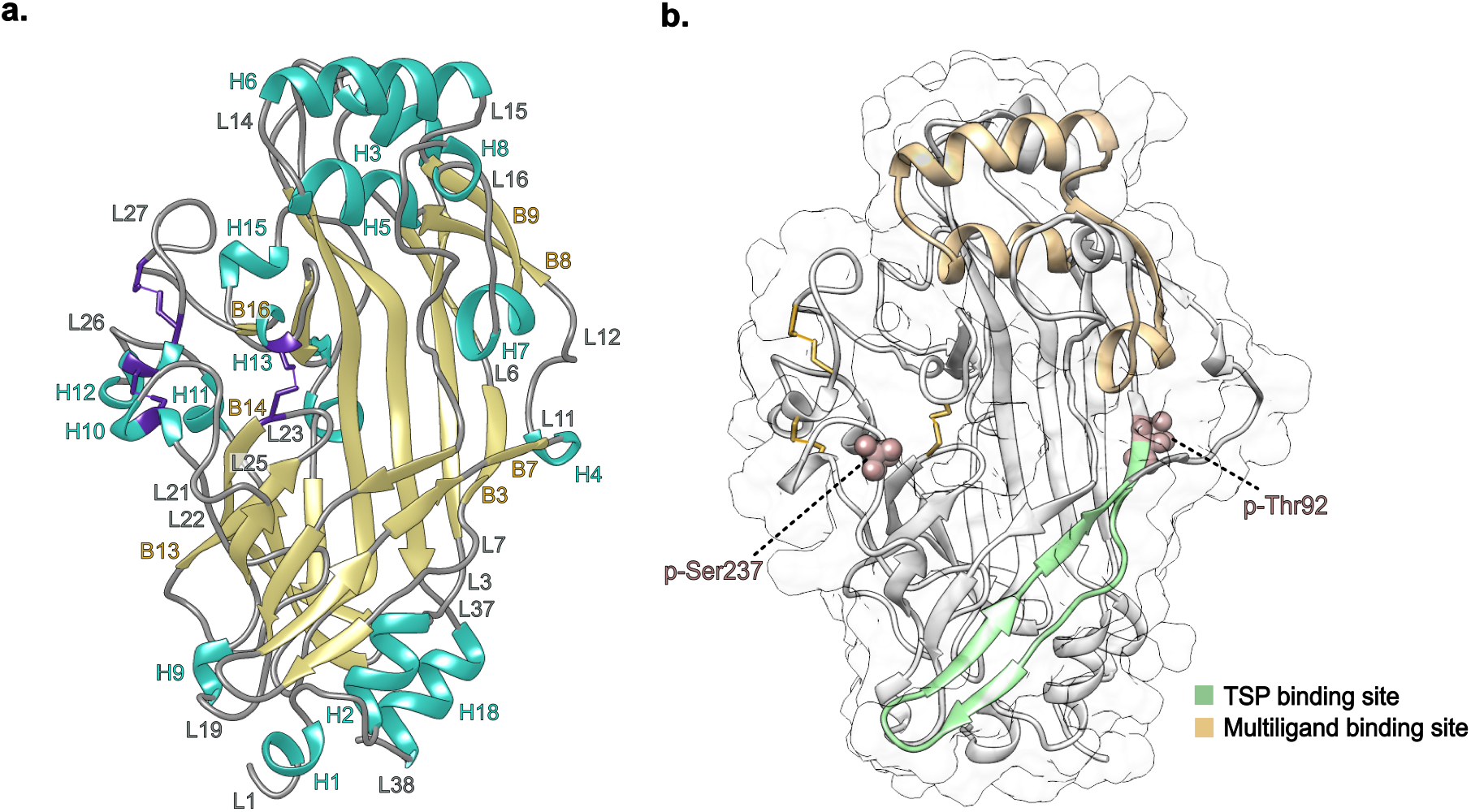
(a) The crystal structure of the CD36 extracellular domain (PDB ID: 5LGD) (Hsieh et al. 2016). Glycosylation sites are removed for clarity. The structure is color coded based on the secondary structure elements, with helices shown in cyan, beta strands in khaki, and loops in gray. Disulfide bonds between residues 243-311, 272-333, and 313-322 are highlighted in purple. The secondary structure elements are numbered (H: helix, L: loop or turn, B: beta strand) by the order they appear in the CD36 sequence. (b) The structure of the CD36 extracellular domain, depicting the thrombospondin-1 (TSP-1) binding site, residues 93-120, in green, and the multiligand binding site, residues 139-183, in khaki. The oxLDL binding site (residues 155-183) overlaps with the multiligand binding site. The phosphorylation sites, Thr92 and Ser237, are represented as spheres.

In human platelets, microvascular endothelial cells, and CD36-transfected fibroblasts, the extracellular domain of CD36 has been reported to be phosphorylated at Thr92, which is determined as a canonical protein kinase C (PKC) consensus site (Asch et al. 1993; Ho et al. 2005). Given the abundance of CD36 in platelets (Nergiz-Unal et al. 2011), most experimental evidence on the effects of phosphorylation on CD36 function has been obtained from platelet studies (Asch et al. 1993; Chu and Silverstein 2012; Guthmann et al. 2002; Hatmi et al. 1996). *In vitro* data demonstrates that phosphorylation of CD36 at Thr92 decreases CD36 binding to TSP-1, with the level of phosphorylation correlating with the inhibition of TSP-1 binding, and markedly reduces its affinity for erythrocytes infected with Plasmodium falciparum (Chu and Silverstein 2012). It has been proposed that the constitutive phosphorylation of CD36 in platelets accounts for the absence of detectable TSP-1 binding in resting platelets, despite high surface expression of CD36. Considering that the interaction between CD36 and TSP-1 inhibits angiogenesis and promotes apoptosis (Klenotic et al. 2013), modulating their binding holds pharmaceutical potential for treating angiogenesis-dependent diseases including cancer (Chignen Possi et al. 2017; Dawson et al. 1999). The presence of the anionic phosphate on Thr92 has been proposed to create steric hindrance that inhibits TSP-1 binding (Klenotic et al. 2013). However, in the absence of atomistic level information on the conformational dynamics of CD36, it remains unclear whether, and to what extent, steric effects govern the interactions between phosphorylated CD36 and TSP-1. It is well documented that phosphorylation can change the energy landscape of the protein (Lätzer, Shen, and Wolynes 2008), and induce conformational changes in the surroundings of the phosphorylation site (Groban, Narayanan, and Jacobson 2006), which can affect ligand binding. The molecular mechanisms by which phosphorylation alters CD36 function remain to be elucidated.

In the plasma membrane, CD36 facilitates fatty acid transport into adipocytes, heart and skeletal muscle cells (Glatz and Luiken 2017). Trafficking of CD36 from endosomes to the plasma membrane regulates the rate of fatty acid uptake (Bonen et al. 2000). Alterations in CD36 mediation of fatty acid metabolism is associated with insulin resistance, diabetic cardiomyopathy, and cardiac hypertrophy (Glatz, Heather, and Luiken 2024). Moreover, cancer cells utilize CD36 to increase fatty acid uptake to meet their metabolic demands to proliferate and metastasize (Feng, Zuppe, and Kurokawa 2023). In platelet cells, phosphorylation of Ser237 leads to a reduction in fatty acid uptake, which is recovered on dephosphorylation (Guthmann et al. 2002). The molecular mechanisms by which CD36 phosphorylation regulates lipid uptake remain poorly understood.

Molecular dynamics (MD) simulations can provide atomic level description of protein structure and dynamics, and sample the conformational changes conferred by post-translational modifications in the absence of experimental data (see review (Weigle, Feng, and Shukla 2022)). MD simulations of various phosphorylated proteins (see examples (Jonniya, Sk, and Kar 2019; Khaled, Gorfe, and Sayyed-Ahmad 2019; Wang et al. 2017; Zhu et al. 2017)), despite being limited to short time scales, demonstrate how phosphorylation alters the conformational dynamics of both the binding site and other distinct loop regions with potential implications for ligand binding. The conformational changes imparted by phosphorylation and other post-translational modifications can however occur on timescales beyond the typical MD simulation sampling time. In a single MD simulation, molecules may be unable to explore all metastable states due to confinement in a local energy minimum. Markov State Models (MSMs) describe the conformational dynamics of a system by discretizing the high-dimensional conformational space into a set of states and by modelling probabilistic, memoryless transitions between the identified states. The transitions between states are stochastic and independent of prior visited states. Therefore, in building MSMs, a single trajectory is not necessarily required to explore the entire phase space, because data from multiple independent simulations can be used to surmount sampling limitations (Bowman, Pande, and Noé 2014).

This study provides an atomistic characterization of the effects of dual phosphorylation (phosphorylation on both Thr92 and Ser237) on the structure and dynamics of the CD36 extracellular domain using MD simulations with a cumulative simulation time of 30 µs. An iterative dimensional reduction protocol identified two regions within CD36 comprising residues 121-131 and residues 296-331, which display slow motions upon phosphorylation. Both regions are located on the protein surface and relatively accessible for protein-protein and protein-ligand interactions. Primarily composed of loop structures, these two regions are inherently flexible and susceptible to rapid dynamics. However, the fact that these regions exhibit the slowest motions within the protein suggests their potential functional significance in the dual phosphorylated form of CD36, as the coordinated slow movement of loops with larger domains can be indicative of allosteric regulation (Skliros et al. 2012). Our findings demonstrate that CD36 phosphorylation induces a conformational rearrangement of the Thr92 adjacent loop (residues 121-131). A cryptic surface pocket emerges near this region upon phosphorylation. Binding site analysis demonstrated that phosphorylation redirects binding of the TSP-1 derived ligands from the canonical TSP-1 binding site (residues 93-120) to the cryptic pocket, providing a potential molecular mechanism for the inhibitory effect of CD36 phosphorylation on TSP-1 binding. Additionally, MSMs indicate that the region spanning residues 296-331, which surrounds one of the two entrances to the internal hydrophobic cavity, undergoes the slowest conformational dynamics in both phosphorylated and unphosphorylated forms, and adopts a more compact and folded conformation upon phosphorylation. Phosphorylation induces partial folding of this region, narrowing the entrance to the internal cavity and reducing its overall volume. Although not always occurring during the simulations nor expected to completely block fatty acid uptake, such narrowing does provide a plausible structural basis for the experimental observations that phosphorylation decreases the rate of fatty acid uptake *in vitro*. We propose that phosphorylation induced changes in CD36 conformational dynamics identified in this study provide a molecular basis for the experimentally observed reductions in TSP-1 binding and fatty acid uptake and should therefore be considered in the development of CD36 targeted antiangiogenic and metabolic therapeutic strategies.

## Results

### Phosphorylation increases the flexibility of a loop region spanning residues 121-131

The structural stability of the phosphorylated and unphosphorylated CD36 was assessed using the root-mean-square deviation (RMSD) as a metric (Figure S1a). Overall, both systems structurally converged within the first 500 ns. RMSD values of CD36 showed no significant difference upon phosphorylation and both systems were stable during the simulations. Phosphorylated CD36 shows more deviation among the replicas in the second half of the simulations. Root-mean-square fluctuation (RMSF) of the C_α_ atoms was used as a metric to examine the structural flexibility. Phosphorylation significantly increased the flexibility of the region spanning residues 121-131 (highlighted in green in Figure S1b).

### Two surface-exposed regions containing loops display the slowest dynamics

Using an iterative dimensionality reduction method based on the time-lagged independent component analysis (tICA) previously described by Barros and coworkers (Barros et al. 2021) (see **Materials and Methods**), we identified two distinct regions of phosphorylated CD36 that contribute most to the slowest motions of the protein. The first region (Reg1) comprises residues 296-331, and spans part of L25, H11, L26, H12, L27, B16, and H13 (Figure 2a, left panel). The second region (Reg2) encompasses residues 121-131, includes B7, L11, H4, and part of L12 (Figure 2a, right panel). The total number of C_α_-C_α_ distances describing the dynamics of these two regions is 38 (Figure 2b), with 21 anchored in Reg1 and 17 in Reg2 (shown as spheres in Figure 2a). The phosphorylated form of CD36 was used as the reference structure to identify the regions with slow dynamics. The unphosphorylated CD36 structure was subjected to the same iterative protocol to determine whether additional regions contributed significantly to its conformational dynamics. Reg1 was the only part of the unphosphorylated CD36 displaying slow dynamics (Table S1), corroborating that the conformational changes observed in Reg2 are induced by phosphorylation.

**Figure 2.**
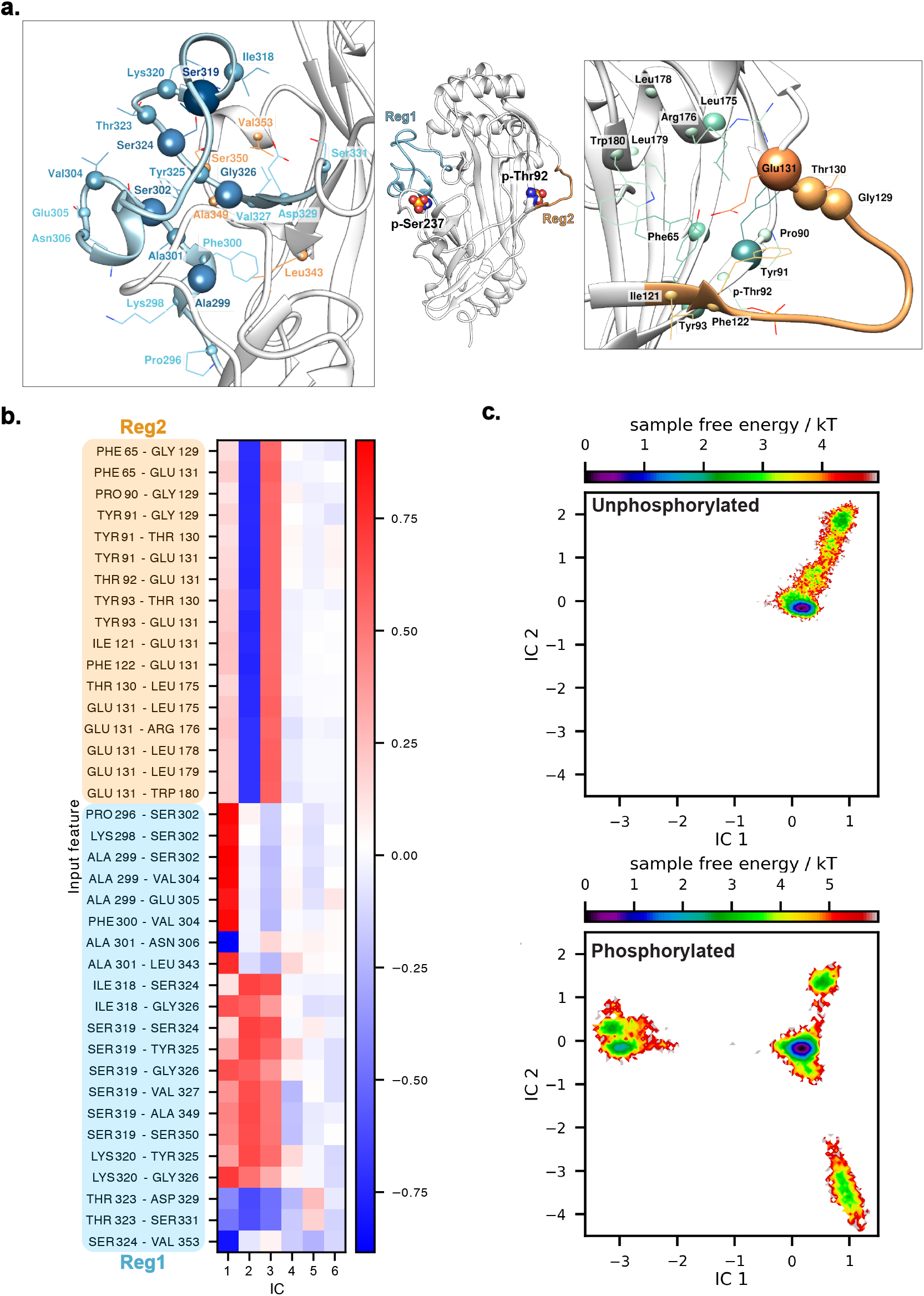
CD36 conformational landscape represented by the selected 38 features. (a) A representative conformation of the phosphorylated CD36 with the phosphorylated residues Thr92 (p-Thr92) and Ser237 (p-Ser237) shown as spheres (middle panel). Blue and orange colored regions indicate Reg1 and Reg2, respectively. Close-up views of both regions are shown (Reg1, left panel; Reg2, right panel), with the C_α_ atoms of the residues used to construct the MSMs represented as spheres. The size of each sphere correlates with the frequency by which a residue appears in the feature list. (b) Input feature correlation with the first six time-lagged independent components (ICs). Residue pairs belonging to Reg1 and Reg2 are highlighted in blue and orange, respectively. (c) Free energy landscape of the unphosphorylated (top) and phosphorylated (bottom) CD36, in terms of the selected 38 features, projected onto the first two independent components.

The first independent components capture the slowest dynamical modes of the protein. The features anchored in Reg1 show a high correlation with the first tICA component (IC1, Figure 2b). Because the tICA components are sorted from slowest to fastest, this indicates that Reg1 is the slowest part of CD36. Reg1-anchored features with a weaker correlation with IC1 were identified in the unphosphorylated CD36 system (Table S1) and were manually added to the feature list to provide a better description of the slow movements in both systems. The C_α_-C_α_ distances anchored in Reg2 are highly correlated with the second tICA component (IC2), demonstrating that this region is the second slowest part of the protein.

The conformational landscape of CD36, represented by the free energy landscape in terms of the first two ICs, changes significantly upon phosphorylation (Figure 2c). For a direct comparison between the two forms of CD36, the conformations explored by each system (represented by the 38 features) were jointly used as an input for tICA, allowing the simulation results of both systems to be mapped into the same low-dimensional coordinate space.

The locations of Thr92 and Ser237 are proximal to Reg2 and Reg1, respectively, within the tertiary structure of CD36. Thr92 was identified in the descriptive feature list of Reg2 in phosphorylated CD36, suggesting that phosphorylated Thr92 directly participates in the dynamics of this region. Neither Ser237 nor its neighbouring residues have been recognized in the feature list describing the slow motions of CD36. To further investigate the dynamics of these regions, separate MSMs were constructed for Reg1 and Reg2 using the identified features.

### The loop comprising residues 121-131 undergoes a closed-to-open motion upon phosphorylation

The conformational landscape of Reg2 (residues 121-131) in terms of the selected 17 features, is shown in Figure 3 for both systems. The conformation of Reg2 remains nearly unchanged within the ensemble of conformers sampled over the 15 µs-long simulation of unphosphorylated CD36 (Figure 3b, Figure S2), which displays only one free energy basin (Figure 3a).

**Figure 3.**
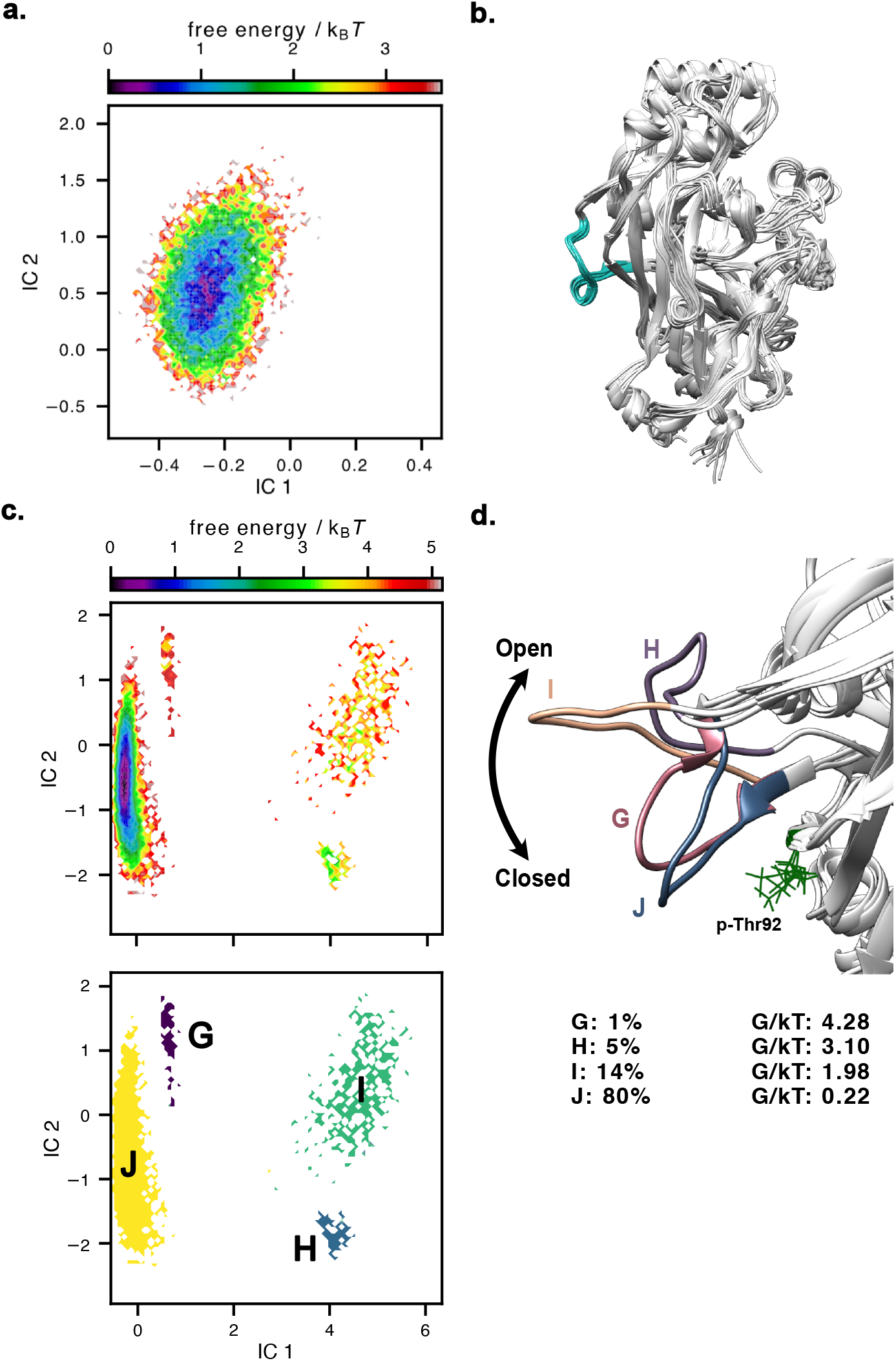
(a) Free energy landscape of Reg2 projected onto the first two independent components for the unphosphorylated CD36. (b) Ten randomly selected representative conformations of the unphosphorylated CD36 from a 15 µs ensemble. Reg2 is shown in cyan. (c) Free energy landscape of Reg2 projected onto the first two independent components (top) and state separation (bottom) in phosphorylated CD36. (d) Representative conformations depicting the metastable states visited by Reg2 upon phosphorylation. The predicted stationary distributions and corresponding free energy are shown. Phosphorylated Thr92 is shown in green.

In contrast, Reg2 undergoes a closed-to-open motion upon phosphorylation and interconverts between two united (four different) metastable states (labeled as G, H, I and J in Figure 3c, d; Figure S3). State J represents the closed state of the Reg2 and displays the highest equilibrium population (Figure 3d). This closed state structurally resembles the conformation of Reg2 observed in the unphosphorylated CD36. To quantify the conformational transition of this loop from closed to open state with respect to Thr92, the average distance between the C_α_ atoms of residues Thr92 and Ser127 (located in the middle of Reg2) is calculated across ten randomly chosen structures. This distance ranges between 9.68 Å in the closed state (state J) to 18.82 Å in state I which represents the conformation in which Reg2 is furthest from Thr92 (Figure S4).

The frequency of transitions between metastable states provides a quantitative understanding of the dynamics of a system. Using the transition path theory (E. and Vanden-Eijnden 2006), PyEMMA calculates the rate of transitions and identifies the pathways between states. The transition rate between two states is inversely proportional to the mean first passage time (MFPT), which indicates the mean time required to transition from one state to another. In phosphorylated CD36, state J (closed state) can be considered as the main conformation for Reg2 to transition between states. The analysis of MFPT between the metastable states of Reg2 (Figure S5) shows that state G is infrequently visited but is separated from state J by the smallest energy barrier. Similarly, the transition from state H to I occurs on a shorter time scale compared to other transitions. In the lower-resolution kinetic behavior, states G and J collectively constitute approximately 81% of the equilibrium population and may be treated as a single state. Similarly, states H and I jointly represent about 19% of the equilibrium conformations.

Visual inspection of the trajectories shows that the conformational changes of Reg2 in the phosphorylated CD36 take place mostly in replicate 1 and the closed-to-open transition of the loop is only observed after 1.5 μs (Figure S6a). Therefore, carrying out short but many simulations may not have revealed this motion. An additional, 2 μs-long simulation seeded from the equilibrated structure in replicate 1 (first frame of the production run) was carried out and incorporated into the existing data, and a new MSM was constructed (Figures S6b, S7). The MSM with the new trajectory has the same number of cluster centers and lag time as the MSM built on the original five trajectories. The new MSM identified three metastable states for Reg2. Due to separate tICA transformations of the data, a direct comparison of the energy landscape between the new MSM and the original one is not possible. Visual inspection of the identified states shows that in the new MSM, states H and I are merged into a single state (state L) with an equilibrium population of approximately 37% (Figure S7b). This aligns with our original findings from the five-trajectory MSM, indicating a small energy barrier between states H and I. State M is the state most similar to unphosphorylated CD36 and shows the highest population as observed in the original MSM. The variance in the stationary properties shows that predicted equilibrium populations may change based on the input data used to construct the MSM, but the identified states remain relatively stable.

The change in the dynamics of Reg2 due to phosphorylation affects the frequency of inter-residue contacts. The most notable differences between phosphorylated and unphosphorylated CD36 were observed in contacts involving the following residues: Tyr62, Gly89, Pro90, Tyr91, Thr92, Ile121, Phe122, Leu126, Ser127, and Val128 (Figure S8) which belong to regions B1, L6, B3, B7, H4, L12 of the crystal structure depicted in Figure 1a. Thr92 interacts with residues Tyr62, Arg63, Gln64, Phe122, Glu123, Leu126, and Ser127 throughout the entire simulation duration in the unphosphorylated state of CD36. These interactions are partially diminished upon phosphorylation, as shown in the contact frequencies outlined in Table S2. Phosphorylated Thr92 makes brief contacts with Pro124, Ser125 and Val128.

### Phosphorylation induces formation of a cryptic pocket on the CD36 surface

Three small pockets surrounding Reg2 were identified in the phosphorylated structure throughout the simulations (Figure 4a). These pockets were transient, appearing fractionally during the simulations. Analysis of the unphosphorylated CD36 demonstrates the formation of pockets 1 and 2. Comparison of pockets 1 and 2 between the unphosphorylated and phosphorylated CD36 indicates that pocket 1 gets formed almost twice as frequently upon phosphorylation (Supplemental Video 1, quantitative data not shown). Pocket 2 is observed in both systems; however, it exhibits a larger volume in the phosphorylated CD36 (Figure S9, Supplemental Video 2).

**Figure 4.**
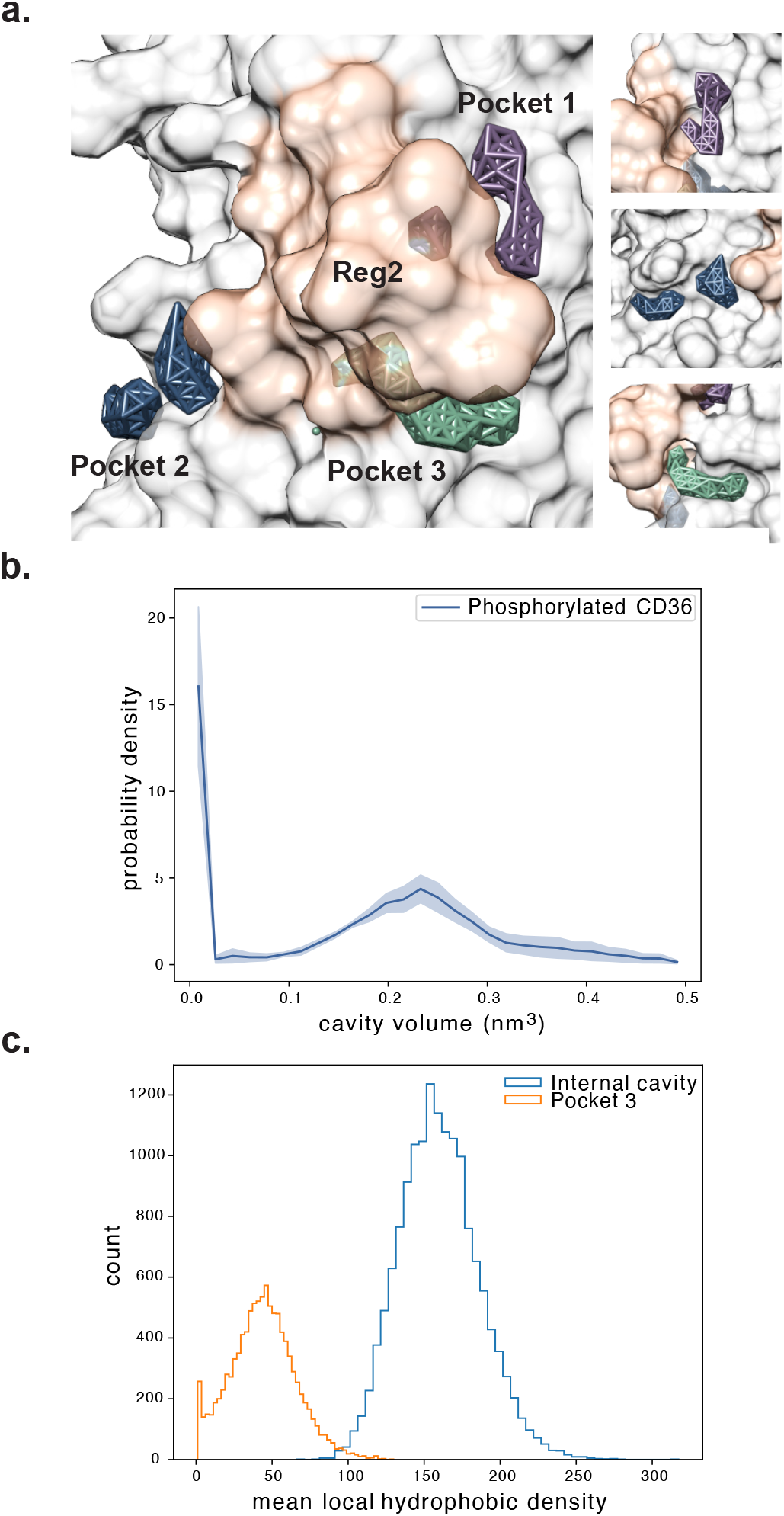
(a) The surface pockets surrounding Reg2 (residues 121-131) shown on the phosphorylated CD36 structure. Reg2 is shown in orange transparent surface representation and pockets 1, 2 and 3 are shown in purple, blue, and green, respectively. (b) Distribution of the volume of pocket 3 averaged over five 3 μs-long trajectories. Shading indicates uncertainty (standard error of the mean) over five independent simulations. (c) Distribution of the mean local hydrophobic density for pocket 3 and the internal cavity of phosphorylated CD36, calculated over 15 µs sampling time. Time frames in which pocket 3 is absent from the structure (∼30% of the 15 µs sampling time) are excluded from the analysis.

Pocket 3 is exclusive to the phosphorylated structure and is located underneath the Reg2 in the three-dimensional structure of the CD36 (Supplemental Video 1). The residues involved in the formation of pocket 3 are listed in Table S3. Regardless of its size, pocket 3 remains present over 70% of the simulation time (15 μs) (Figure 4b, Figure S10). To evaluate the potential of pocket 3 as a ligand binding site, its mean local hydrophobic density was calculated. This metric quantifies the spatial fraction of hydrophobic regions within a pocket where hydrophobic residues are most accessible to hydrophobic drug like molecules (Schmidtke et al. 2011; Schmidtke and Barril 2010). Comparison of this metric with that of the internal cavity of CD36, which is the primary hydrophobic cavity responsible for fatty acid uptake (Hsieh et al. 2016), indicates that pocket 3 possesses favorable druggable characteristics despite being significantly smaller than the internal cavity (Figure 4c). These findings suggest that pocket 3 may serve as a binding site for modulators designed to selectively inhibit phosphorylated CD36.

### Phosphorylation alters the thrombospondin binding site of CD36

The CLESH domain of CD36, encompassing residues 93–120, serves as the structural recognition site for TSP type I repeats (TSRs) (Chu and Silverstein 2012; Crombie and Silverstein 1998). The minimal TSP-1 region required for CD36 binding is the TSR2 domain, in which experimentally identified key residues include Trp420, Trp423, Trp426, Ile438, Arg440, Arg442, and Lys464. Examination of mutant constructs by NMR spectroscopy and migration assays has shown that the positively charged surface formed by the Cys, Trp, Arg (CWR)-layered core of TSR2 interacts with the negatively charged acidic residues Glu101, Asp106, Glu108, and Asp109 of the CD36 CLESH domain (Klenotic et al. 2011, 2013).

Dual phosphorylation of CD36 does not measurably affect the structure and dynamics of the neighbouring CLESH domain. An initial attempt to construct an MSM on the CLESH domain did not yield clearly separated metastable states, likely because the domain undergoes subtle conformational fluctuations that do not produce sufficiently distinct or well-separated conformational states for reliable state assignment. Principal component analysis (PCA) followed by k-means clustering, performed using PyEMMA on the positions of CLESH C_α_ atoms, identified two conformational populations that are highly similar (Figure S11a-c). The backbone RMSD between the two structures corresponding to their respective cluster-centers was 1.0 Å, further supporting the absence of a major conformational rearrangements in this region throughout the simulations. Comparison of the phosphorylated and unphosphorylated CD36 revealed no notable differences in the solvent accessible surface area (SASA) of the CLESH domain, suggesting that its solvent exposure remained unaffected by phosphorylation (Figure S11d).

Collectively, the data presented above support the hypothesis that phosphorylation-induced conformational changes in Reg2 and the formation of a cryptic surface pocket near the CLESH domain modulate TSP-1 binding to CD36. This hypothesis was evaluated by assessing the binding of two sets of peptides derived from the TSR2 domain of TSP-1 (Figure 5). The first set consists of two ligands containing different subsets of the key TSR2 residues experimentally identified as contributing to CD36 binding (Figure 5a, b). Ligand 1 spans residues 418-443 of TSR2, encompassing the CWR-layered core and all experimentally identified CLESH binding residues except Lys464. Ligand 2 (residues 438-464 of TSR2) contains Ile438, Arg440, Arg442, and Lys464, but lacks the CWR-layered core. Both ligands have the same charge (+2) but differ in hydrophilicity (Table S4). The second set comprises three clinically or preclinically tested peptides developed by the Abbott Laboratories based on the parent TSR2 sequence GVITRIR, named ABT- 526, ABT-510 (Haviv et al. 2005), and ABT-898 (Garside et al. 2010) (Figure 5c). All peptides in the ABT series are capped with acetyl and ethylamide groups at the N and C-termini, respectively. Compared to the parent TSR2 antiangiogenic sequence, GVITRIR, ABT-526 has two additional residues at the termini, sarcosine (Sar) at the N terminus, and proline at the C-terminus. It features a d-isoleucine (d-Ile) in place of isoleucine, and the internal arginine in the parent peptide is replaced with a neutral non-canonical amino acid, norvaline (Nva). ABT-510 incorporates d-allo-isoleucine (d-allo-Ile) substitution in place of the d-isoleucine present in ABT-526, resulting in significantly improved solubility and longer half-life (Haviv et al. 2005), which supported its advancement into clinical trials. ABT-898 omits the initial sarcosine residue present in ABT-526 and ABT-510, and features modifications in the central sequence, namely serine in place of threonine, and glutamine in place of norvaline. For all ligands, binding site searches were performed within the region spanning residues 92-131 of CD36, which encompasses both the CLESH domain and Reg2, to guide Surflex-dock (Jain 2003) for protomol generation (see **Materials and Methods**). When target residues are used for protomol generation in Surflex-dock, they define the intended binding site for protomol-guided docking rather than a strict geometric boundary on ligand placement (Lohning et al. 2017). The protomol captures the local site surface through hydrophobic and polar probes, and Surflex-dock searches this space by aligning ligand fragments to the protomol, rebuilding the full ligand, and refining poses through local and all-atom optimization (Chaput and Mouawad 2017; Jain 2003, 2007, 2009). Since docked poses are scored against the receptor rather than the protomol alone, the final ligand pose can extend into neighboring pocket regions if those interactions are more favorable (Holt, Chaires, and Trent 2008; Jain 2007; Lohning et al. 2017). Following protomol generation, ensemble docking was carried out using 20 CD36 conformations each for phosphorylated and unphosphorylated form (see **Materials and Methods** for the rationale for selecting these conformations).

**Figure 5.**
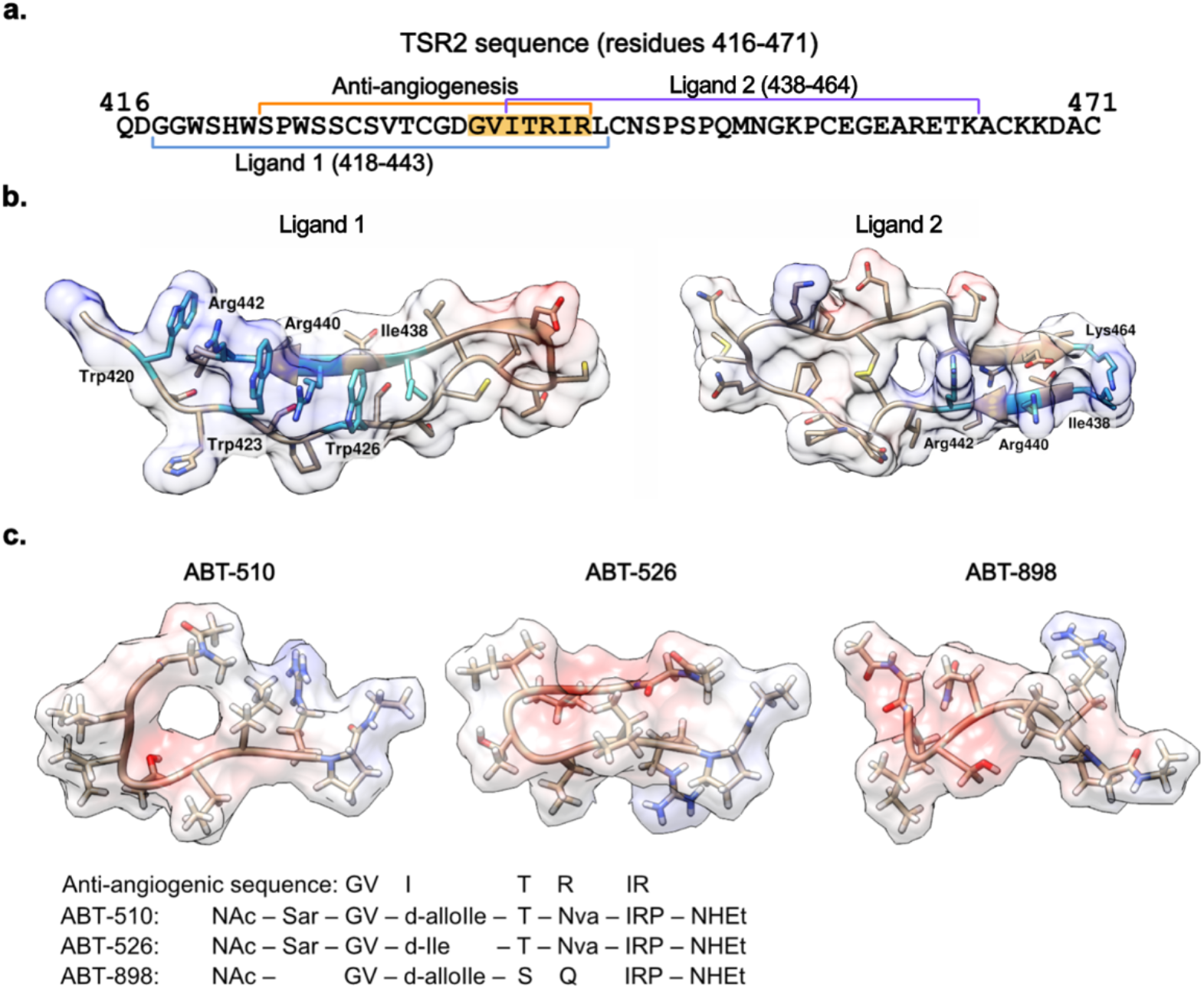
(a) Sequence of the TSR2 domain of TSP-1. Within TSR2, a 19-amino-acid segment, SPWSSASVTAGDGVITRIR, marked with the orange line, was experimentally identified to have anti-angiogenic activity. From this segment, a 7-residue part, GVITRIR, highlighted in yellow, serves as the foundation for the design of the ABT series of TSP-1 mimetic peptides with anti-angiogenic properties. Peptide ligands 1 and 2 span residues 418-443 and 438-464 of TSR2, respectively. (b) Structures of the peptide ligands 1 and 2. Residues that are experimentally identified to contribute to binding the CD36 CLESH domain are highlighted in cyan and labeled. (c) Structures of ABT-510, ABT-526, and ABT-898 peptides, derived from the parent TSR2 sequence, GVITRIR. The Coulombic surface coloring method in UCSF Chimera was used with the color scale ranging from -10 (red) to 0 (white) to 10 (blue) in units of kcal/mol·e at 298K.

Binding assessment of the first set of peptides demonstrates that phosphorylation shifts the binding site of TSP-1-derived peptides towards the cryptic surface pocket adjacent to Reg2. Ligand 1 binds to the unphosphorylated CD36 through electrostatic interactions between its positively charged CWR-layered core and the negatively charged Asp106, Glu108, and Asp109 residues of the CLESH domain (Figure 6b, Table S5). This binding mode is in agreement with the previously reported experimental findings (Klenotic et al. 2011, 2013), and serves as a validation of the docking method used. It occurs in 8 out of 10 docking models examined using different sets of randomly selected structures of the unphosphorylated CD36. In the remaining two docking positions that are less energetically favorable, binding is driven predominantly by hydrophobic contacts rather than electrostatic interactions, with the binding site remaining near the CLESH domain but relocating closer to Reg2 (Figure S13). Ligand 2 binds to the unphosphorylated CD36 through electrostatic interactions with Glu101, Asp106, and Glu108 of the CLESH domain. The change in the peptide sequence and hydrophilicity (5 positive and 3 negatively charged residues) enhances the electrostatic interactions between CD36 and ligand 2 (Figure S14b, Table S5).

**Figure 6.**
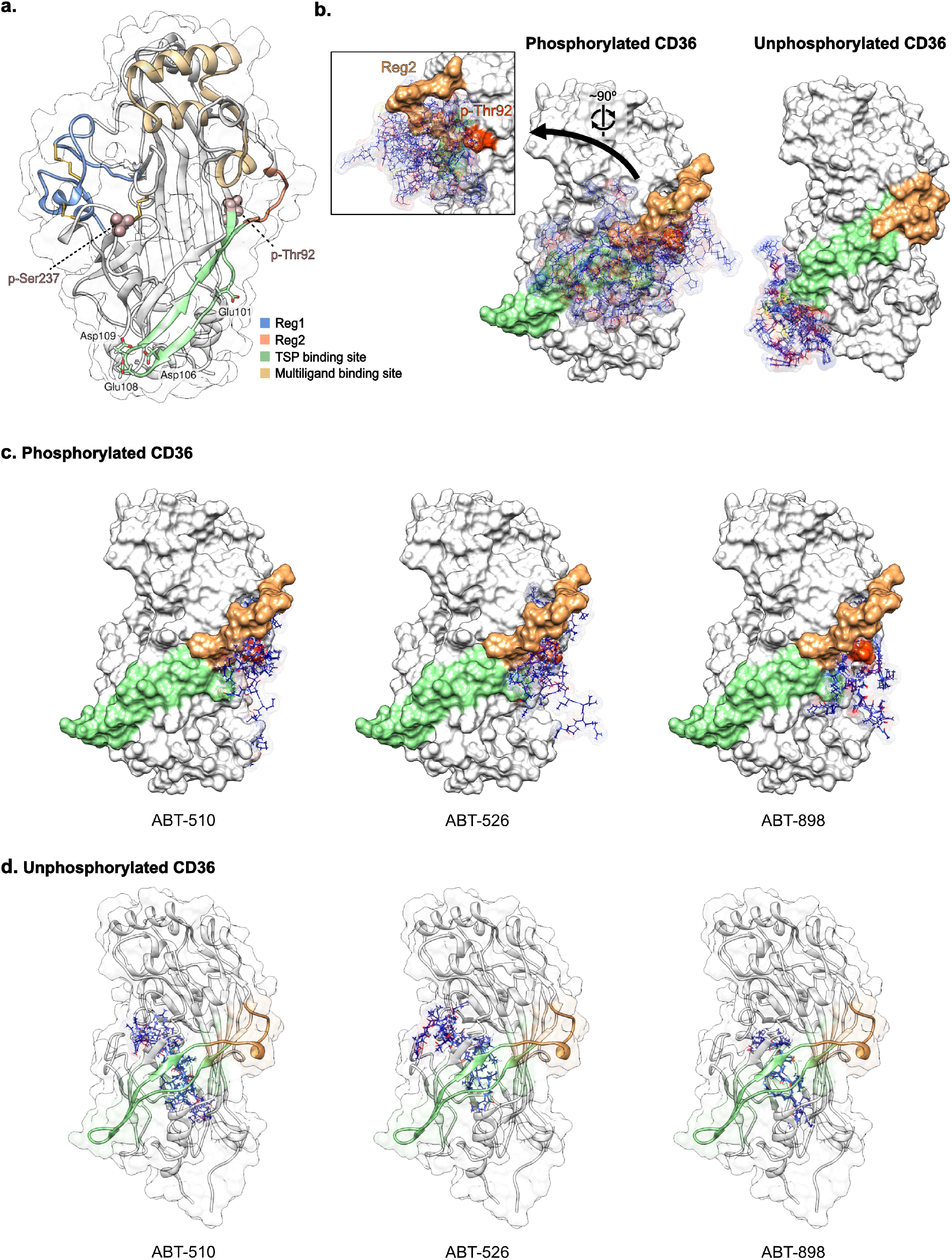
(a) A random frame from MD trajectories showing the extracellular domain of phosphorylated human CD36. The TSP binding site (CLESH domain spanning residues 93-120) and multiligand binding site (residues 139-183) are depicted in green and khaki, respectively. The oxLDL binding site (residues 155-183) overlaps with the multiligand binding site. The phosphorylation sites, Thr92 and Ser237, are represented as spheres. Two regions showing slow movements, identified as Reg1 (residues 296-331) and Reg2 (residues 121-131), are highlighted in blue and orange, respectively. The negatively charged acidic residues Glu101, Asp106, Glu108, Asp109 in the TSP binding site are specified. (b) Docking of peptide 1 derived from TSP-1 type 1 repeat 2 (TSR2) (residues 418-443) to phosphorylated and unphosphorylated CD36. CD36 structure is shown in solid surface representation; the top five ligand binding poses are depicted in wire representation. The p-Thr92 is highlighted in red. A closeup from a 90° rotation shows the top five ligand binding poses of phosphorylated CD36 with respect to Reg2 and Thr92. (c) Docking of the ABT peptides to (c) phosphorylated and (d) unphosphorylated CD36. Top five binding poses of the peptide ligands ABT-510, ABT-526, and ABT-898 are shown in wire representation.

Independent of sequence differences, both ligands 1 and 2 bind the phosphorylated CD36 about Reg2, demonstrating the relevance of this site for TSR2 affinity (Figure 6b, Figure S14a). Phosphorylation-induced conformational changes reduce the accessibility of acidic residues in the CLESH domain. Of the acidic residues that mediate electrostatic interactions with ligands in unphosphorylated CD36, only Glu101 retains its contact with ligand 1 in phosphorylated CD36 (Table S5). In the phosphorylated CD36, the loss of these interactions is compensated by the hydrophobic and electrostatic interactions formed by Reg2, p-Thr92, and the neighboring residues (Table S5).

Ensemble docking of the ABT peptide series revealed a phosphorylation-dependent shift in binding site comparable to that observed for the TSR2-derived ligands 1 and 2. In the unphosphorylated CD36, ABT-510, ABT-526, and ABT-898 docked favorably within the internal hydrophobic cavity, away from Reg2 and without apparent contacts with the exposed CLESH surface (Figure 6d). This binding mode differs from that of the TSR2 derived ligands. Previous experimental studies have mapped the TSR2 binding interface on CD36 to the CLESH domain (Klenotic et al. 2013), but have not established a corresponding binding interface for the ABT peptides. ABT peptides have been reported to exhibit functional profiles different from that of the parent TSR-derived peptide. For example, ABT-510 has weaker effects on fatty acid uptake and nitric oxide signaling but induces caspace activation more strongly than the parent peptide, suggesting that ABT peptides may engage CD36 differently or elicit different downstream signaling responses (Isenberg et al. 2008). Thus, previous studies support CD36 binding by ABT peptides without requiring them to reproduce the full canonical TSR2 binding surface (Audet et al. 2013; Dawson et al. 1997). Upon phosphorylation, all three ABT peptides redirect their binding to the newly formed cryptic surface pocket near Reg2, mirroring the behavior of ligand 1 and ligand 2 (Figure 6c), further supporting the hypothesis that this phosphorylation-induced cryptic pocket acts as the primary binding determinant for the TSP-1 mimetic peptides on phosphorylated CD36.

Experimental studies on a recombinant extended CLESH construct (residues 81 to 120 of CD36) suggest that TSR2 binds this extended domain, and that phosphorylation of Thr92 sterically inhibits this interaction (Chu and Silverstein 2012; Klenotic et al. 2013). However, these studies are limited to the boundaries of the CD36 construct tested and therefore cannot determine whether phosphorylation induces alternative TSR2 binding sites beyond those boundaries. Our simulations demonstrated that phosphorylation affects the conformational dynamics of the residues 121-131 of CD36 (Reg2) and opens a cryptic pocket at an adjacent region. Although study of the entire complex between CD36 and TSP-1 is challenging due to the approximately 2.5 times larger size of TSP-1 compared to CD36, such simulations could reveal other binding interactions beyond those of TSR2 which were demonstrated in our analysis.

### Phosphorylation increases the helicity of a helix-loop region close to Ser237

The conformational landscape of Reg1 (the helix-loop region comprising residues 296 to 331), in terms of the selected 21 features (see **Materials and Methods**), is projected onto the first two independent components (Figure 7). tICA transformation of the feature data using a lag time of *τ* =10 ns reduced the dimensionality from 21 to 14, capturing 95% of the total dynamic variance of the protein. The MSMs identified four metastable states in both systems, corresponding to the free energy minima of the sampled conformations (Figure 7). States A, B, C, and D represent the four metastable states visited by the unphosphorylated CD36. States C and D of the unphosphorylated CD36 are retained in the phosphorylated form of CD36 with a significant decrease in their stationary populations; 35% to 4% for state C, 39% to 9% for state D. Upon phosphorylation, Reg1 of CD36 sample two new conformations (states E and F) that were not detected in the unphosphorylated form, together accounting for ∼87% of the equilibrium population (Figure 8, Figure S15). In both states E and F, Reg1 adopts a more helical structure. On average, the number of residues showing helical content in the unphosphorylated CD36 is ∼9, which increases to ∼12 upon phosphorylation (Figure S16). The observed increase in helicity following serine phosphorylation is consistent with previous studies (Andrew et al. 2002; Rani and Mallajosyula 2021; Signarvic and DeGrado 2003; Smart and McCammon 1999), which have shown that the extent of this structural change depends on the position of the phosphorylated serine within the protein structure. Additionally, phosphorylation stabilizes the transient conformations of the protein, as evidenced by the significant increase in the timescale of transitions between different states, with the phosphorylated CD36 displaying MFPTs more than 200 *μ*s (Figure S17b) while the highest MFPT observed in the unphosphorylated CD36 is 5.5 *μ*s (Figure S17a).

**Figure 7.**
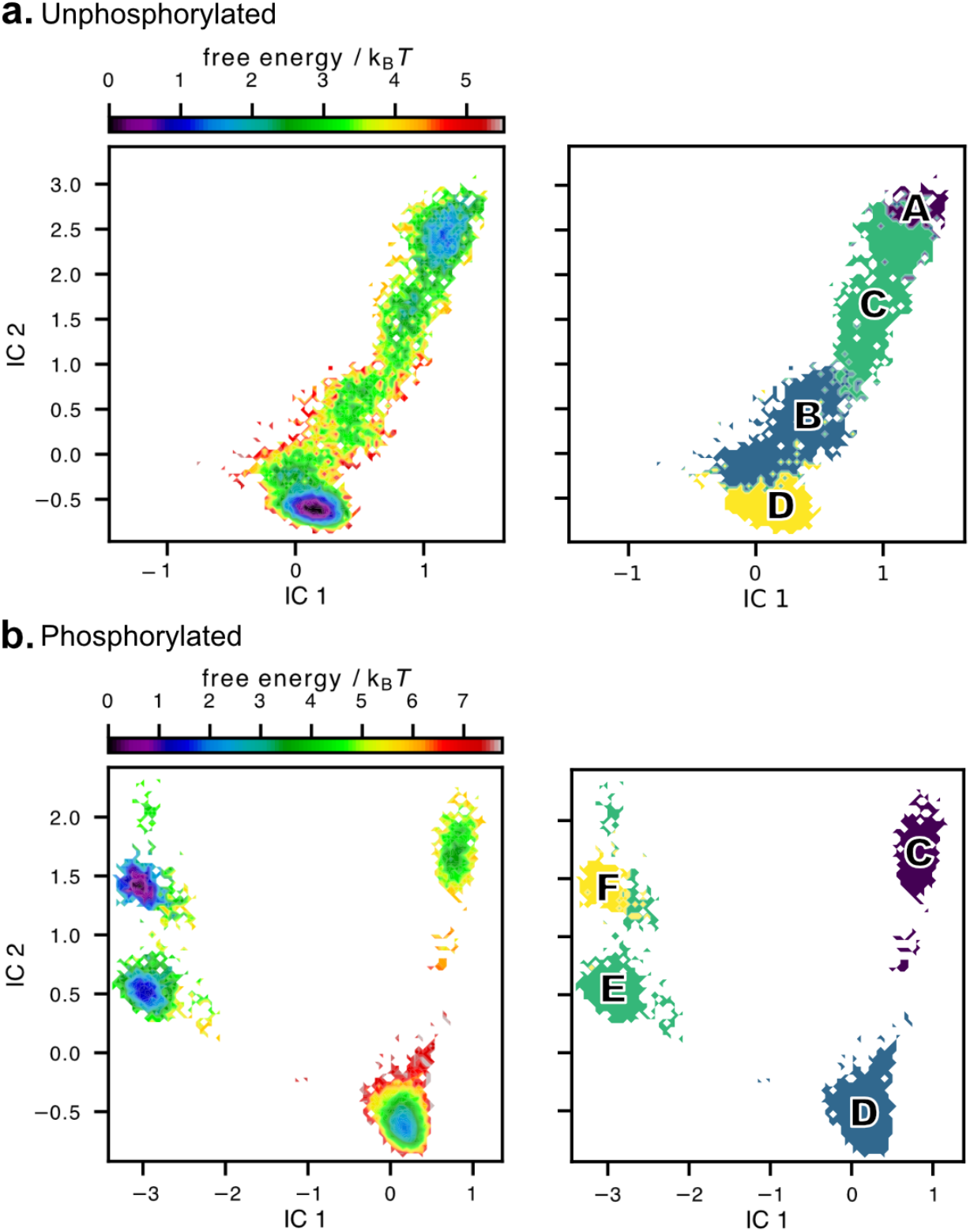
Free energy landscape of the helix-loop region comprising residues 296 to 331 (left) and the corresponding metastable states (right) projected onto the first two independent components for the (a) unphosphorylated and (b) phosphorylated CD36. The unphosphorylated CD36 system exhibits four metastable states labeled as A, B, C, and D. Upon phosphorylation, states C and D remain, while two new metastable states, labeled E and F, emerge in the system.

**Figure 8.**
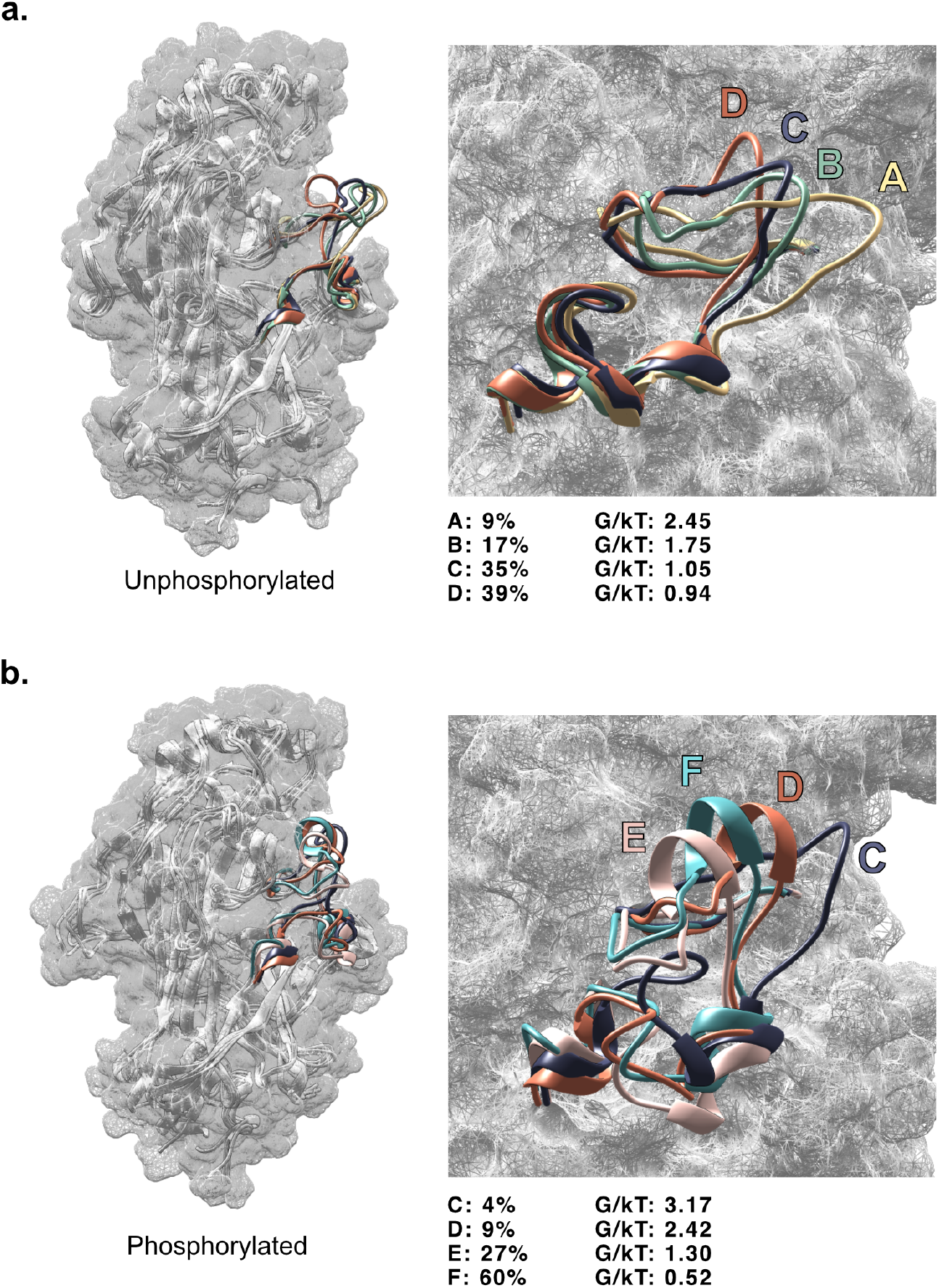
Representative conformations for each metastable state of the helix-loop region comprising residues 296 to 331, randomly selected from a 15 µs ensemble for (a) unphosphorylated and (b) phosphorylated CD36 systems. The stationary distributions and the corresponding free energies predicted by MSMs for each state are displayed.

### Phosphorylation limits access to the internal hydrophobic cavity and decreases its effective volume

The crystal structure of CD36 (PDB ID: 5LGD) comprises two long-chain fatty acids in its hydrophobic internal cavity (Hsieh et al. 2016). These fatty acids were removed from the structure prior to the simulations presented in this study. There are two entrances to the internal cavity which are the likely entry points for fatty acids (Figure S18a). Reg1, which is spatially close to Ser237, encompasses one of the entrances (Entrance 2 in Figure S18a). While experimental data indicate that phosphorylation of Ser237 impairs fatty acid uptake in platelet cells (Guthmann et al. 2002; Shu et al. 2022), the molecular basis for this effect has yet to be elucidated. We hypothesize that conformational changes induced by phosphorylation may either limit access to the internal cavity or reduce its effective volume, making it less favorable for fatty acid transport. Given the spatial proximity of Ser237 to Entrance 2, the addition of a bulky phosphate group may obstruct access to the internal cavity. To test this hypothesis, the size of the opening of Entrance 2 was estimated by measuring the distances between the α-carbons of the central Ile318 residue in Reg1 and those of Asp352 and Asn408 at the mouth of Entrance 2. Phosphorylation of CD36 was found to narrow the joint probability distributions of these distances indicating a shrinking of the opening of Entrance 2 (Figure 9a, b). Additionally, the volume of the internal cavity was estimated and compared throughout the MD trajectories of the phosphorylated and unphosphorylated CD36. On average, the distribution of the mean cavity volume shows that the interior hydrophobic cavity becomes smaller upon phosphorylation, with a decrease from 5.81 nm^3^ (unphosphorylated) to 4.89 nm^3^ upon phosphorylation (Figure 9c). This decrease of 920 Å³ in volume corresponds to roughly twice the crystallographic molecular volume of a 16-carbon long palmitic acid molecule, estimated as approximately 414.5 Å³ per molecule from the unit cell parameters (V = 1658.0 Å³, Z = 4) (Moreno et al. 2006). Collectively, the reduced accessibility of the internal cavity and the decrease in its effective volume may explain the previously reported reduction in the rate of fatty acid uptake in platelets following CD36 phosphorylation (Guthmann et al. 2002).

**Figure 9.**
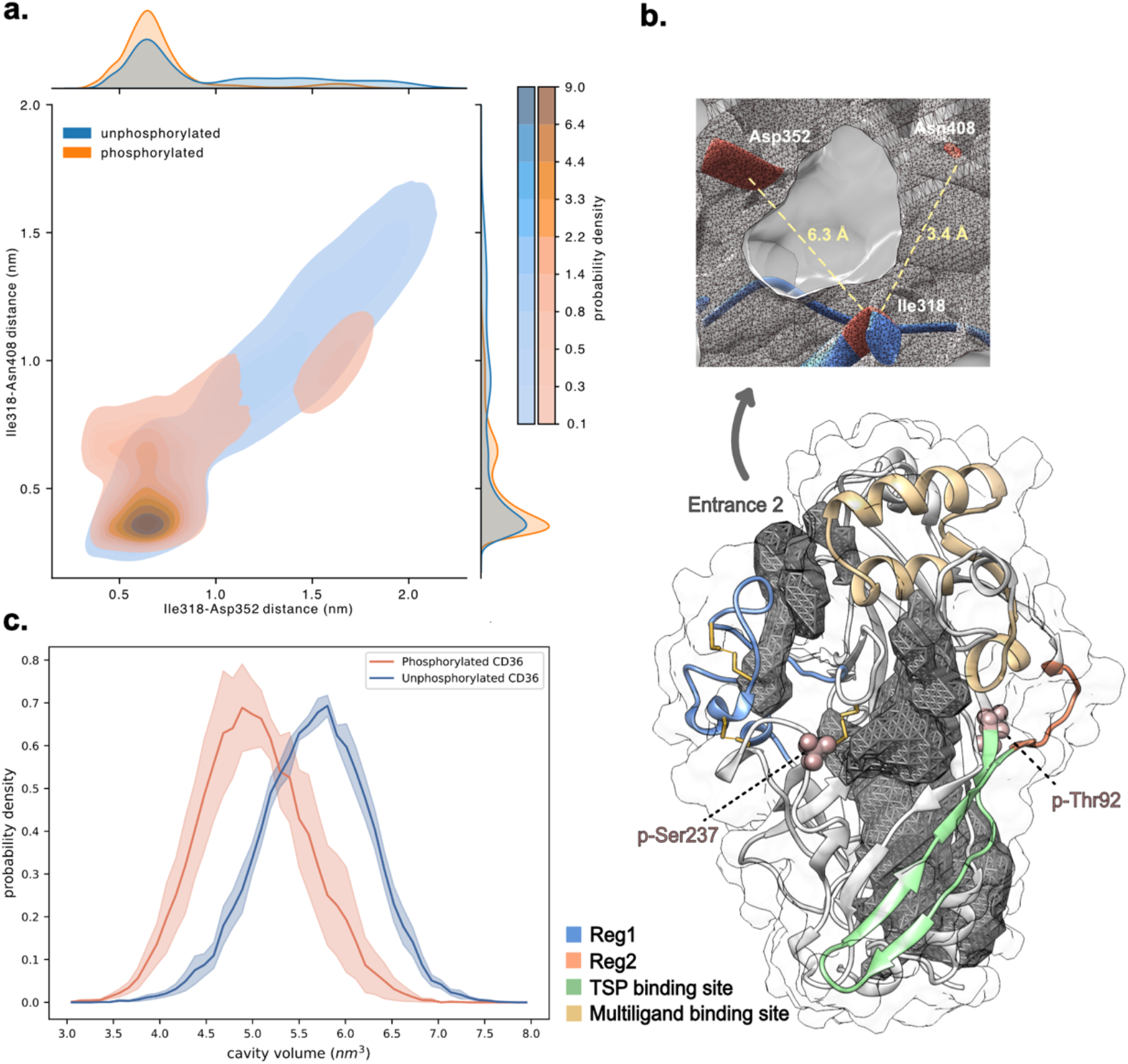
(a) Joint probability distribution of the distance between Ile318-Asp352 and Ile318-Asn408 over the entire ensemble of unphosphorylated (in blue) and phosphorylated (in orange) CD36. In both phosphorylated and unphosphorylated CD36, the most probable distance from Ile318 to Asp352 and Ile318 to Asn408 is observed at 0.63 nm and 0.34 nm, respectively. (b) A random frame selected from the MD trajectories of the dual phosphorylated CD36. The thrombospondin and multiligand binding sites are depicted in green and khaki, respectively. The oxLDL binding site (residues 155-183) overlaps with the multiligand binding site. Residues Thr92 and Ser237 are represented as spheres. Two regions showing slow movements, identified as Reg1 (residues 296-331) and Reg2 (residues 121-131), are highlighted in blue and orange, respectively. The internal hydrophobic cavities of CD36 are shown in gray. A closeup of Entrance 2 is shown on the top with the most probable distance between Ile318 from Asp352 and Asn408 highlighted. (c) Probability distribution of internal hydrophobic cavity volume of CD36 calculated over 15 μs ensemble for unphosphorylated (in blue) and phosphorylated (in orange) CD36. Shading indicates uncertainty (standard error of the mean) over five independent simulations.

## Discussion

Phosphorylation is one of the most prevalent post translational modifications. It regulates protein – protein interactions and protein activity by inducing structural and dynamic changes either locally at the phosphorylation site or allosterically at distal regions of the protein (Subhadarshini et al. 2024). Experimental evidence supports phosphorylation of more than two-thirds of the human proteome, while proteomics-based estimates suggest that up to 90% of proteins may undergo phosphorylation. Despite their prevalence and importance in signaling, phosphoproteins remain underrepresented in PDB with only a small fraction of phosphosites having solved structures (Correa Marrero et al. 2025).

The crystal structure of CD36 extracellular domain was resolved in an unphosphorylated state (Hsieh et al. 2016), despite experimental evidence supporting its phosphorylation (Asch et al. 1993). Structural characterization of phosphoproteins remains challenging and requires a combination of complementary experimental and computational biophysical methods. Phosphoproteins often exist as a heterogeneous mixture of unphosphorylated, partially or fully phosphorylated states, which can compromise structural resolution. Many phosphorylation sites are located within flexible loops or intrinsically disordered regions, precluding crystallization. Additionally, phosphorylation typically induces subtle shifts in conformational ensembles rather than large-scale changes in the protein fold (Correa Marrero et al. 2025) requiring conformational analysis at the ensemble level. A systematic characterization of CD36 phosphorylation under basal conditions across diverse cell and tissue types is lacking and is essential to elucidate the tissue specific, functional roles of extracellular domain phosphorylation.

Phosphorylation of Thr92 in the CD36 extracellular domain inhibits TSP-1 binding in a manner proportional to the extent of phosphorylation (Chu and Silverstein 2012). The CD36 – TSP axis mediates angiogenesis, inflammation and coagulation across multiple diseases, including cancer, vascular and metabolic disorders. In cancer, interactions between CD36 and TSP-1 can elicit both tumor suppressive and tumor promoting effects. In the tumor microenvironment, binding of TSP- 1 to CD36 can activate different downstream signalling events with antiangiogenic and tumor suppressive outcomes, which include apoptosis of endothelial cells, suppression of vascular endothelial growth factor (VEGF) driven cell migration and tube formation, as well as inhibition of nitric oxide signalling and myristic acid uptake (Isenberg et al. 2008). Conversely, CD36 – TSP- 1 interactions have also been reported to promote metastasis in prostate cancer (Firlej et al. 2011) and leukemia (Farge et al. 2023), with these effects partly attributed to TSP-1 mediated increases in hypoxia. Beyond cancer, downregulation of TSP-1 and the associated reduction in CD36 mediated inhibition of angiogenesis contributes to choroidal neovascularization in ocular diseases such as wet age-related macular degeneration (Choi et al. 2023) and diabetic retinopathy (Puchałowicz and Rać 2020). In the absence of a comprehensive characterization of CD36 phosphorylation across disease contexts, the extent to which phosphorylation modulates CD36 – TSP-1 binding and influences diseases-promoting or disease-suppressive outcomes remains unclear.

Treatment of diseases that are driven by excessive formation of new blood vessels, such as cancer, diabetic retinopathy, and wet age-related macular degeneration, requires restoring a healthy angiogenic balance. Utilizing the antiangiogenic endothelial TSP-1 – CD36 interactions, Abbott Laboratories developed a series of peptides based on the parent TSR2 sequence GVITRIR, namely ABT-526, ABT-510 (Haviv et al. 2005), and ABT-898 (Garside et al. 2010) (Figure 5c), and tested their preclinical or clinical efficacy. In preclinical models of mouse and canine cancer, ABT-526 caused a drastic reduction in microvascular density and inhibited tumor growth (Isenberg et al. 2008). Phase II trials of ABT-510 in advanced renal cell carcinoma (Ebbinghaus et al. 2007) and metastatic melanoma (Markovic et al. 2007) demonstrated that the drug was well tolerated but exhibited limited efficacy as a monotherapy, ultimately leading to the discontinuation of its clinical development. Preclinical studies on ABT-898 demonstrated an improved half-life and decreased tumor growth in animal models of ovarian cancer (Campbell et al. 2011) and soft tissue sarcoma (Sahora et al. 2012). When formulated as a protein-like polymer, ABT-898 successfully inhibited angiogenesis in mouse models of wet AMD (Choi et al. 2023). Despite the efficacy in animal models, ABT-898 did not advance to human trials and is mainly used as a veterinary drug. Our results provide a plausible molecular basis likely contributing to the insufficient clinical efficacy of TSP based mimetics, including the ABT series. We present the first evidence, to our knowledge, for a cryptic pocket on the surface of the dual phosphorylated CD36 extracellular domain, that is absent in the unphosphorylated state but is present only in the phosphorylated state. This pocket exhibits high affinity for TSR2 and redirects the binding of TSR2 mimetics away from the canonical CD36 TSR-2 binding region, the CLESH domain, which is responsible for initiating antiangiogenic signaling. To overcome this phosphorylation induced “drug resistance”, potential strategies could involve designing and co-delivering molecules that selectively occupy the cryptic pocket, thereby redirecting TSP-1 mimetics to the canonical CD36 CLESH binding site and restoring the initiation of antiangiogenic signaling.

CD36 facilitates the cellular uptake of long-chain fatty acids and mediates lipid signaling in the heart, skeletal muscle, adipose tissue, liver, and immune cells (Glatz and Luiken 2018; Li et al. 2025). Dysregulation of CD36 mediated lipid handling contributes to the development and progression of a broad spectrum of cardiometabolic diseases, metabolic disorders, and multiple cancers (Chen et al. 2022; Yang et al. 2024). CD36 overexpression in cancer cells enhances the uptake of extracellular fatty acids, thereby promoting tumor growth and metastasis (Feng et al. 2023). *In vitro* data show that phosphorylation of the CD36 extracellular domain by protein kinase A (PKA), a kinase known to phosphorylate Ser237, significantly decreases palmitate uptake by intact human platelets, with uptake restored in the absence of phosphorylation (Guthmann et al. 2002). This is further supported by *in vitro* studies in small intestinal cells demonstrating that dephosphorylation of CD36 enhances or restores fatty acid uptake (Lynes et al. 2011). Conversely, PKA activation was reported not to affect CD36 mediated fatty acid uptake in isolated rat cardiac myocytes, suggesting that cardiac muscle may lack this ectokinase mediated regulatory pathway of CD36. Thus, CD36 phosphorylation may have limited functional relevance in cardiac muscle and in diseases involving this tissue (Luiken et al. 2002). Our simulations provide a mechanistic insight into how phosphorylation alters the structure and dynamics of the CD36 extracellular domain and how these changes may influence its fatty acid uptake. Our data demonstrate that phosphorylation narrows one of the openings to the CD36 internal cavity and reduces its overall effective volume, with a reduction corresponding to approximately the volume occupied by two 16-carbon long palmitic acid molecules. This structural contraction is dynamic and provides a plausible molecular mechanism for the phosphorylation dependent reduction in fatty acid uptake reported *in vitro* (Guthmann et al. 2002). Our simulations report on the dual phosphorylated form of CD36 and therefore do not isolate the specific effects of phosphorylation at Ser237. However, given the closer proximity of Ser237 to the internal cavity opening, which becomes narrower upon phosphorylation, we propose that phosphorylation at this site is the primary contributor to the observed structural effects.

## Conclusions

The present study provides an atomistic characterization of how dual phosphorylation influences the structure and dynamics of CD36 and proposes molecular mechanisms underlying the phosphorylation-mediated reduction in TSP-1 binding and fatty acid uptake. Our data demonstrate that phosphorylation (1) significantly alters the dynamics of a surface exposed loop adjacent to the CLESH domain (residues 121-131); (2) opens a cryptic surface pocket with high affinity for TSR- derived peptide ligands; (3) increases the helicity of residues 296-311, thereby narrowing one of the entrances to the internal cavity; and (4) reduces the volume of the internal cavity. Ensemble docking of both longer TSP-1 derived ligands and smaller clinically and preclinically tested peptides demonstrate the high binding affinity of the cryptic pocket, which forms near phosphorylated Thr92. To our knowledge, this is the first evidence that phosphorylation can open a cryptic pocket on the CD36 surface. We propose that the shift in ligand binding away from the canonical CLESH domain provides a molecular mechanism underlying the experimentally observed reduction in TSP-1 binding upon CD36 phosphorylation and, consequently, the attenuation of antiangiogenic signaling. Furthermore, the phosphorylation dependent decrease in the internal cavity volume corresponds approximately to the combined molecular volume of two 16-carbon palmitic acid molecules, matching the number and size of the lipids resolved in the crystal structure of the CD36 extracellular domain. We propose that the reduced accessibility and volume of the internal cavity upon phosphorylation collectively provide a mechanistic explanation for the previously reported reduction in fatty acid uptake by platelets following CD36 phosphorylation.

## Materials and Methods

### Simulation Details

The crystal structure of the human CD36 extracellular domain comprising residues 35 to 434 was obtained from the chain A of the Protein Data Bank (PDB) entry 5LGD (Hsieh et al. 2016). The pdb file of CD36 was edited by removing the crystallographic water molecules, glycans and fatty acids present in the crystal structure. To phosphorylate the residues Thr92 and Ser237, the PDB Reader & Manipulator tool of the CHARMM GUI online server (CHARMM-GUI n.d.) was used. The phosphorylated forms of both residues were modelled as di-anionic phosphates (PO_4_^2-^). All-atom classical molecular dynamics simulations were carried out using GROMACS version 2021.2 (GROMACS 2021.2 Manual n.d.) with the CHARMM36m force field (April 2015). The parameters for the phosphorylated residues were assigned by analogy (Table S6). Three disulfide bonds were created between Cys243-Cys311, Cys272-Cys333, and Cys313-Cys322. For each system, five independent simulations were carried out. Periodic boundary conditions were applied in all directions. The systems were solvated using TIP3P water in a dodecahedron box. To obtain a salt concentration of 0.15 M, Na^+^ and Cl^−^ ions were added. The N and C termini were charged and set to NH_3_^+^ and COO^−^, respectively. Energy minimization was carried out using the steepest descent algorithm. Electrostatic forces were treated using Particle-Mesh Ewald (PME) and the cut- off for long-range interactions was set to 1.2 nm. Virtual sites were used for hydrogen atoms, and a 4-fs time step was used for the simulations. To relax the CD36 structure gradually, a multi-step position restraint protocol was used, reducing the force constant for position restraints on heavy atoms from 1000 kJ.mol^-1^.nm^-2^ to 0 kJ.mol^-1^.nm^-2^ in 11 steps. The force constants used in these steps were 1000, 500, 250, 120, 60, 30, 15, 10, 5, 1, 0 kJ.mol^-1^.nm^-2^. The position restraint simulations lasted 220 ns in total. The system was maintained at a temperature of 298 K and a pressure of 1 bar. The GROMACS V-rescale (Bussi, Donadio, and Parrinello 2007) method was used for temperature coupling, which rescales the kinetic energy of the system by randomly selecting the target kinetic energy from the canonical equilibrium distribution. NPT equilibration was carried out for 20 ns, with pressure control performed in two stages: initially using the Berenden method for 10 ns, followed by the Parrinello-Rahman algorithm for the remaining 10 ns. Production simulations were carried out for 3 µs, totalling 15 µs for the unphosphorylated CD36 and 15 µs for the phosphorylated CD36. Root-mean-square deviation (RMSD), root-mean-square fluctuation (RMSF), solvent accessible surface area (SASA) and secondary structure calculations were performed using the built-in tools of GROMACS, *gmx rms*, *gmx rmsf*, *gmx sasa* and *gmx do_dssp*, respectively.

### Markov State Model Construction

To investigate how phosphorylation affects the conformational ensembles of CD36, MSMs were constructed using the Python-based software PyEMMA, version 2.5.7 (Scherer et al. 2015). The trajectories were thinned by saving data at every 1 ns. Five independent trajectories were used to construct the MSM for each system.

The first step in building MSMs is to choose a feature set that describes the slow dynamics of the system. A commonly used method for feature selection is to intuitively define three or four feature sets and calculate the variational approach for Markov processes (VAMP) scores for each (Mardt et al. 2018; McGibbon and Pande 2015; Wu and Noé 2020). This method becomes challenging in the case of larger proteins due to the high-dimensional nature of the feature arrays. In this study, the pairwise distances between C_α_ atoms were selected as the feature set, as previously reported as an optimal feature for describing protein dynamics (Scherer et al. 2015). The feature data was refined using an iterative method based on the time-lagged independent component analysis (tICA) (Pérez-Hernández et al. 2013). This approach was originally introduced by Barros and coworkers (Barros et al. 2021) and further explored by Koulgi and coworkers (Koulgi et al. 2022).

### Feature data refinement

The initial feature dataset contained all possible C_α_-C_α_ distances in the structure of the CD36 extracellular domain, totalling 79,003 features. Initially, the distances exceeding 10 Å and those less than 3 Å throughout all simulations, were removed from the dataset. Subsequently, data with a variance smaller than 0.005 was eliminated and pairs involving terminal residues were excluded, which resulted in 1,372 features. Finally, the iterative tICA-based method was applied to this data set, and data showing low correlation with either of the first two components (ICs) were removed. To do this, in every step of this method, tICA transformation within PyEMMA was used to linearly transform the feature data to a lower dimensional space containing 95% of the cumulative kinetic variance. tICA identifies data points with the highest autocorrelation at a given lag time, enabling the detection of the slower components within the dataset. Following this transformation, feature data with correlation values below a predefined cut- off was excluded from further analysis. The cut-off values for the correlation coefficients in the iterative analysis were set to 0.4, 0.6, 0.7, 0.8, 0.85, 0.9. Both positive and negative correlations were considered. This multi-step data refinement protocol resulted in 38 features describing the most significant dynamics of CD36 structure. The details of these steps are given in Table S7.

### MSM construction

Out of the initially selected 38 pairs, 21 pairs were anchored in the region containing residues 296-331 (Reg1). The remaining 17 pairs were linked to the region with residues 121-131 (Reg2) (Figure 2b). Reg1 includes part of L25, H11, L26, H12, L27, B16, and part of H13, while Reg2 comprises part of B7, L11, H4, and part of L12 (Figure 1a). Separate MSMs were constructed for Reg1 and Reg2 using the selected 21 and 17 C_α_-C_α_ distances, respectively. The projection of the concatenated trajectories onto the first four tICA components for Reg1 (Figure S19) and Reg2 (Figure S6) show that metastability is reasonably described. The independent trajectories show variation in the frequency of the discreet jumps in each component, suggesting that some of the slow motions are only sampled in a subset of trajectories.

For each region, the feature data from the phosphorylated and unphosphorylated CD36 systems were concatenated, and this joint data was subjected to a tICA transformation, resulting in a shared low-dimensional space for both systems. This approach ensures a direct comparison between the two systems. A lag time of 10 ns for Reg1 and 5 ns for Reg2 were chosen in the tICA transformation, reducing the dimensionality from 21 to 14 in Reg1 and from 17 to 9 in Reg2. Subsequently, the transformed data for the phosphorylated and unphosphorylated CD36 systems were divided into separate sets to construct MSMs for each system individually.

In the next step, trajectories of the transformed feature data were clustered into a set of microstates using k-means clustering. This step is necessary to determine the transition matrix between the microstates. 75 and 30 cluster centers were chosen for Reg1 and Reg2, respectively. The appropriate lag time for all the MSMs was determined by examining the implied timescale (ITS) plots. Lag times of 10 ns and 5 ns were selected for Reg1 (Figure S20a, b) and Reg2 (Figure S21a), respectively. Chapman−Kolmogorov test was used to validate the MSMs (Figure S20c, d; Figure S21b). According to Chapman−Kolmogorov, when the system’s dynamics follow Markovian principles, progressing through an MSM for *n* steps with a lag time of *τ* should be equivalent to using an MSM with a single lag time of *nτ*, as shown in Eqn. (1).

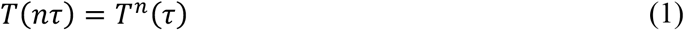

In Eqn. (1), *T* is the transition probability matrix. The implied time scale or relaxation time scale is calculated using Eqn. (2):

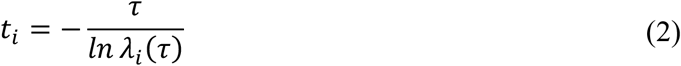

in which *λ_i_* (*τ*) is an eigenvalue of *T*(*τ*) (Bowman et al. 2014). When the Chapman-Kolmogorov equation remains valid for the MSM, multiplying the lag time in Eqn. (2) by an integer value *n* will not alter the value of the implied time scale. For this reason, picking the lag time from the ITS plot when leveled off ensures that the selected lag time maintains Markovian behaviour. After determining the number of metastable states from the spectral analysis of the MSMs (Figure S22), Robust Perron Cluster Analysis (PCCA+) (Röblitz and Weber 2013) algorithm within PyEMMA was used to assign the microstates to the identified metastable states (Figure S23).

Upon visual examination of the trajectories, the majority of conformational changes in Reg2 of the phosphorylated CD36 was observed to occur in the second half of one of the five replicates (replicate 1). To further investigate the accessible conformational space of the phosphorylated CD36, an additional simulation was seeded from the first frame of the production run in replicate 1 and carried out for 2 μs. This additional trajectory was combined with the existing five trajectories, and a new MSM was constructed using the same set of 17 C_α_-C_α_ distances for Reg2. Results from the analysis of this MSM are provided in Figure S7.

### Analysis of the internal cavity and pockets

MDpocket (version 4.0) (Schmidtke et al. 2011) was used with default parameters to examine the dynamic behavior of the internal hydrophobic cavity and surface binding pockets. To isolate the internal cavity, the *-S* flag was used during pocket detection, which excludes the surface-exposed, small pockets. To obtain a good estimate of the volume and extent of the cavity, an iso-value of 0.1 was chosen to extract the frequency grid points. For analysis of smaller surface pockets, the iso-value was set to 1.5 to extract density grid points, allowing for a more focused examination of these specific pockets.

### Ensemble docking

Ensemble docking was carried out to investigate the binding of the peptides derived from the TSP- 1 type I repeat 2 (TSR2) to the CD36 extracellular domain. The crystal structure of TSR2 (PDB ID: 1LSL (Tan et al. 2002)) was truncated to comprise respectively residues 418 to 443 and 438 to 464 as peptide ligands 1 and 2 (Figure 5a,b). Due to the limitations imposed by the docking software the entire TSR2 was impossible to use as a single ligand. The second set of TSR2-derived peptides, ABT-510, ABT-526, and ABT-898, were constructed using RosettaCommons (Leaver-Fay et al. 2011) and subjected to 250 ns MD simulations. The final frame from each simulation was used as the ligand structure for docking (Figure 5c). An ensemble of 20 CD36 conformations was generated from the MD simulation trajectories and the TSR2 peptide ligands were docked to this ensemble, using the BioPharmics Platform docking module, Surflex-dock (version 5.189) (Jain 2003). For the phosphorylated CD36, five representative structures from each metastable state identified for residues 121-131 were used for the docking. For the unphosphorylated CD36, 20 random conformers were selected from the simulation trajectories, because Reg2 does not undergo any significant conformational change throughout the simulations of the unphosphorylated system. For binding site definition, residues 92-131 of CD36, which include both the CLESH domain and Reg2, were specified to guide Surflex-dock for “protomol” generation. A protomol is a pseudo-molecule, serving as an idealized active site ligand used as a target to generate potential ligand binding poses (Jain 2003).

### Other analyses

To investigate the influences of phosphorylation on the inter-residue interactions, the frequency of the contacts formed across the entire structure was calculated for both phosphorylated and unphosphorylated CD36 systems, using the Contact Map Explorer Python package (Contact Map Explorer — Contact Map Explorer 0.7.1.dev0 documentation n.d.), an extension of MDTraj (McGibbon et al. 2015). The cut-off for the residue contacts was the default value of 0.45 nm. VMD (version 1.9.4a55) and UCSF Chimera (version 1.17.1) were used to visualize the structures and trajectories.

## Supplementary Material Description

The supplementary material includes additional figures and tables supporting the main findings. Figures S1–S23 illustrate analyses of the conformational dynamics, structural flexibility, contact frequency differences, metastable state representations, solvent accessibility, pocket volume distributions, and ensemble docking results for phosphorylated and unphosphorylated CD36 systems. Tables S1–S6 provide detailed data on identified features for MSM construction, residue contacts, ligand hydropathicity, docking interface residues, force field parameters for phosphorylation, and data processing for MSM construction. Together, these materials offer a comprehensive view of the methods, models, and molecular insights discussed in the main text.

## Data availability

The source data and the code used for analysis can be found at: https://doi.org/10.5281/zenodo.22002585

## Acknowledgments

GG and DME acknowledge computational resources provided by Calcul Québec (www.calculquebec.ca) and the Alliance de recherche numérique du Canada (https://alliancecan.ca). DME acknowledges funding support by CIHR Commercialization Project Grant (PJT-186296) and NSERC Discovery Grant (RGPIN-2022-05185), and equipment support by Canada Foundation for Innovation John R. Evans Leaders Fund.

