## Supporting Information for "CD36 phosphorylation alters the thrombospondin binding site and reduces internal cavity accessibility and volume"

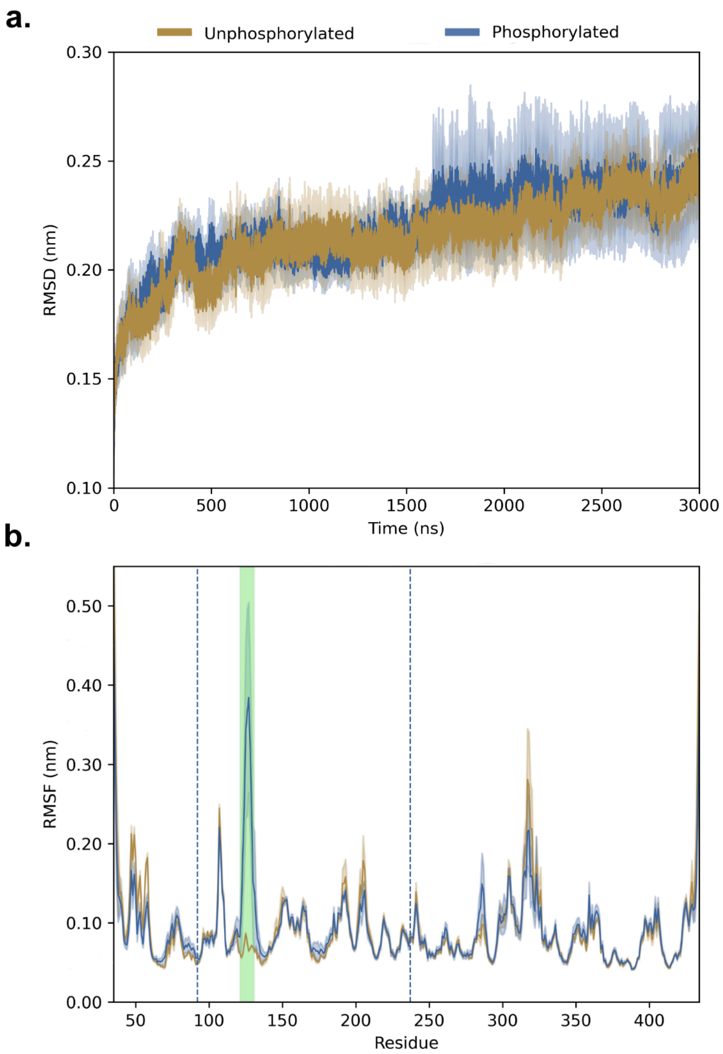


**Figure S1.** Stability and flexibility of the phosphorylated and unphosphorylated forms of CD36. (a) Root-mean-square deviation (RMSD) of the heavy atoms of CD36 relative to the first frame of the production simulation. (b) Root-mean-square fluctuation (RMSF) of the Cα atoms of CD36. Phosphorylation sites (Thr92 and Ser237) are shown with dashed lines. The residues 121-131 (highlighted in green) show significantly higher flexibility in the phosphorylated CD36 structure compared to the unphosphorylated CD36. Shading indicates uncertainty (standard error of the mean) over five independent simulations.

**Table S1.** The list of features identified in the unphosphorylated CD36 system. All the identified features are anchored in Reg1.

| | |
|---|---|
| ILE 318 - SER 324 | SER 319 - ALA 349 |
| ILE 318 - GLY 326 | SER 319 - SER 350 |
| SER 319 - SER 324 | LYS 320 - TYR 325 |
| SER 319 - TYR 325 | LYS 320 - GLY 326 |
| SER 319 - GLY 326 | THR 323 - ASP 329 |
| SER 319 - VAL 327 | THR 323 - SER 331 |

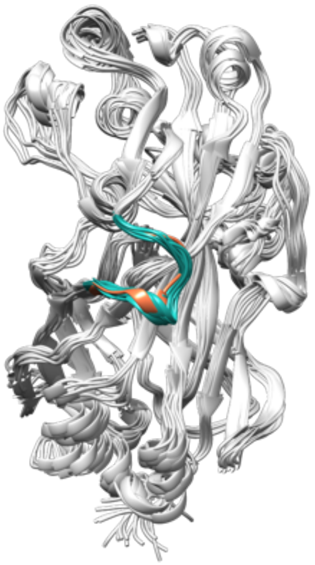

**Figure S2.** 10 randomly selected representative conformations for unphosphorylated CD36 from a 15 $\mu$s ensemble (cyan) overlaid on top of the crystal structure of CD36 (PDB ID: 5LGD) (orange).

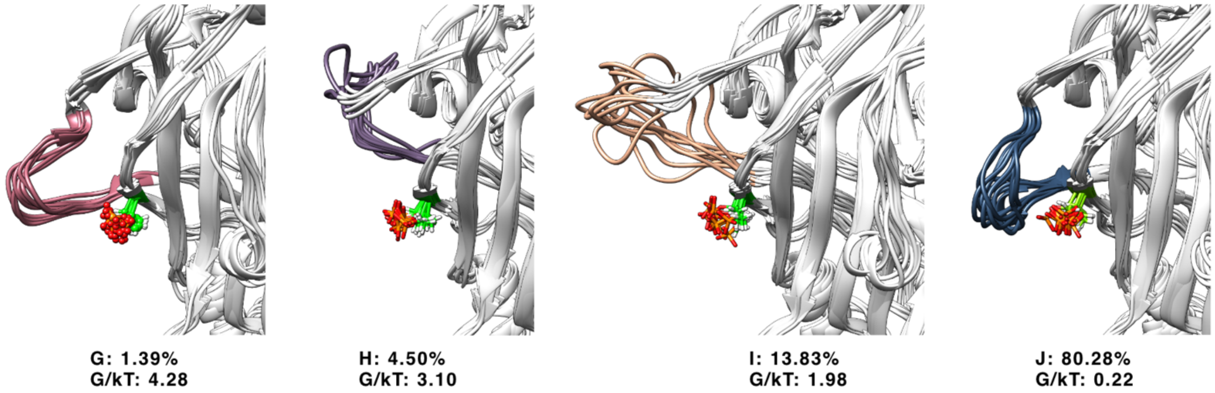


**Figure S3.** Ten randomly selected representative conformations of Reg2 from each metastable state identified from the 15-μs ensemble of phosphorylated CD36. p-Thr92 is shown as a ball-and-stick model colored by heteroatoms.

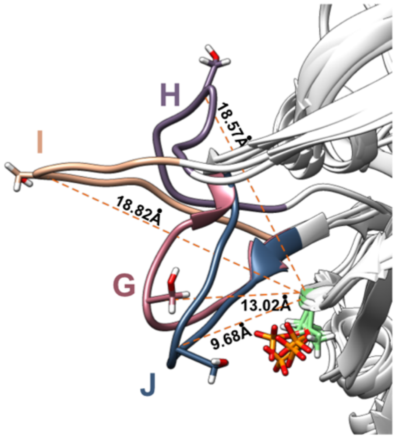


**Figure S4.** Representative conformations of the four metastable states visited by Reg2 in phosphorylated CD36. The average distance between the $C_\alpha$ atoms of Thr92 and Ser127, calculated over 10 randomly selected conformers, for each state is shown.

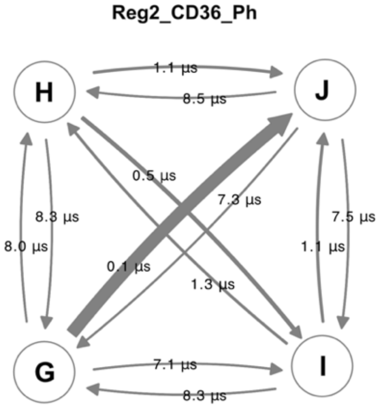


**Figure S5.** Kinetic model showing the mean first passage time (MFPT) across states for Reg2 in phosphorylated CD36.

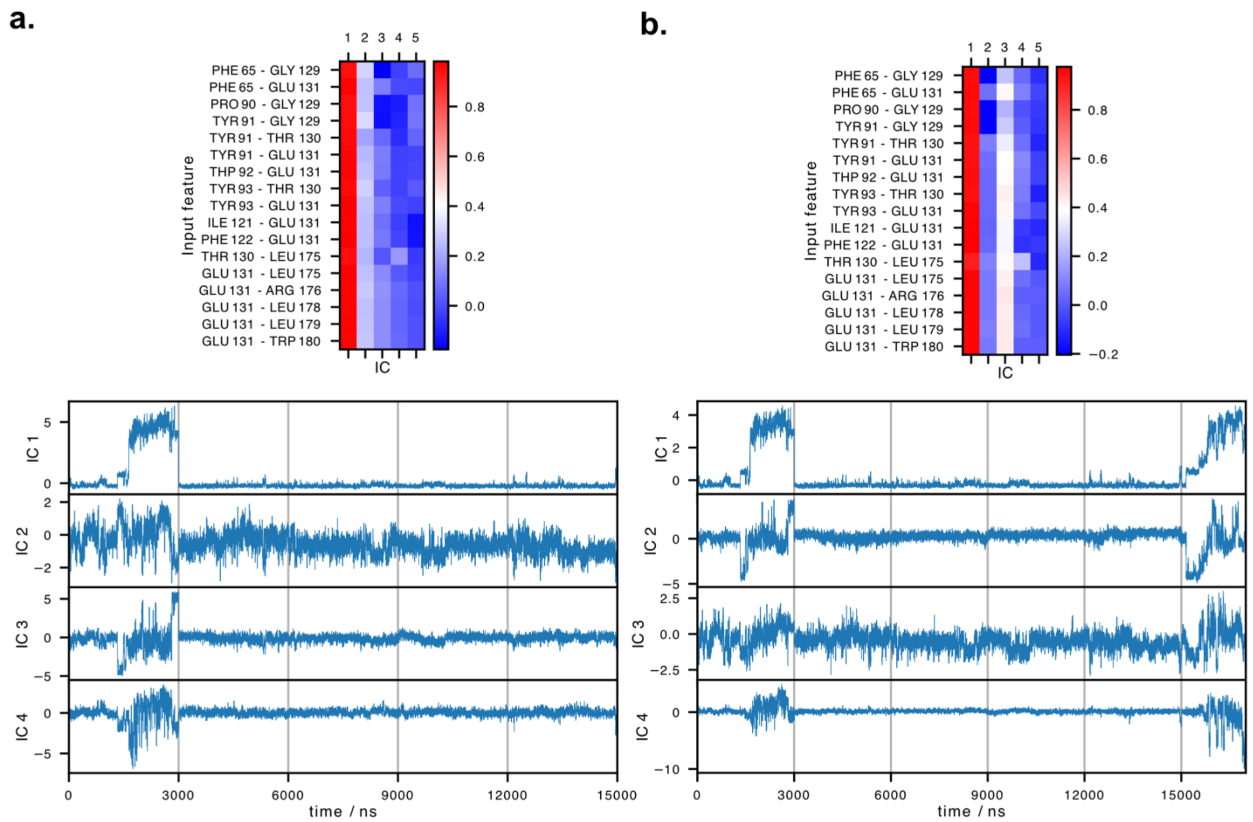


**Figure S6.** Projection of the concatenated phosphorylated CD36 trajectories onto the top TICA components: (a) the original five trajectories and (b) the original five trajectories together with the sixth additional trajectory. The top panels show the correlations between the 17 selected features and the first five independent components (ICs) (lag time=5). The bottom panels show all trajectories projected onto the first four ICs. Individual trajectories are separated by gray lines.

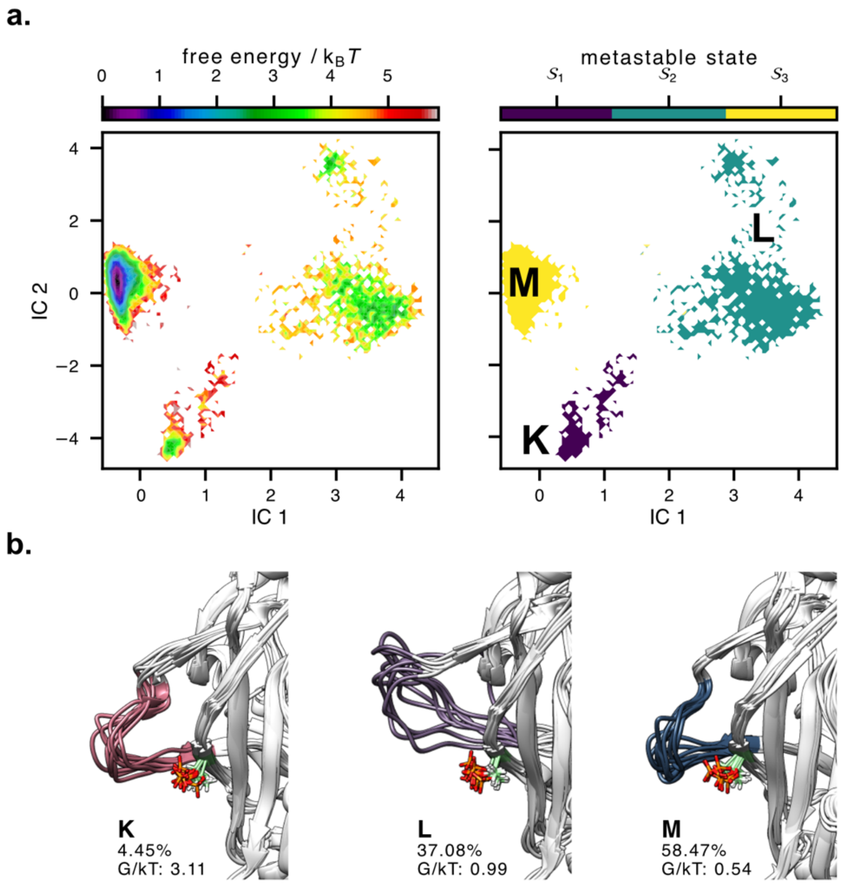


**Figure S7.** Updated MSM results incorporating the sixth trajectory for Reg2 in phosphorylated CD36. (a) Free energy landscape projected on the first two ICs (left) and corresponding metastable states (right). (b) Ten randomly selected representative conformations for each metastable state of Reg2 from the 17 μs ensemble. p-Thr92 is shown as licorice model.

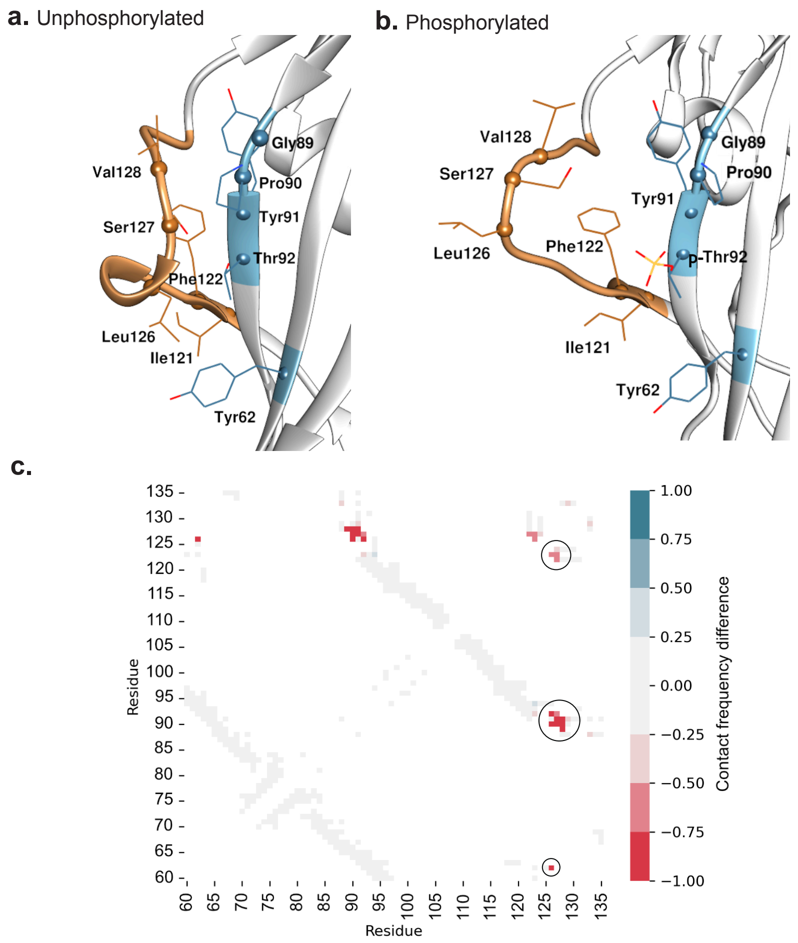


**Figure S8.** Residues displaying the highest difference in contact frequency between (a) unphosphorylated and (b) phosphorylated CD36, with Reg2 highlighted in orange and the $C_\alpha$ atoms of the specified residues shown as spheres. (c) Difference in the frequency of residue contacts between the phosphorylated and unphosphorylated CD36. The map shows the contact frequencies of the unphosphorylated system subtracted from those of the phosphorylated system. The regions highlighted in red correspond to the contacts which exist exclusively in the unphosphorylated CD36. The largest differences are circled indicating contacts made by residues Leu126, Ser127, and Val128 from Reg2. Specifically, Leu126 interacts with residues Tyr62, Pro90, Thr92, and Phe122. Ser127 forms contacts with Pro90, Tyr91, Thr92, Ile121, and Phe122. Val128 interacts with Gly89, Pro90, and Tyr91.

**Table S2.** Most common contacts made by residues Thr92 and Ser237 in unphosphorylated and phosphorylated CD36 throughout the simulations. The contact frequency values are shown for each residue. Asp244 and Lys233 remain as the interacting residues with Ser237, with an observed increase in their contact frequency in the phosphorylated CD36. In the unphosphorylated state of CD36, Thr92 consistently interacts with Tyr62, Arg63, Gln64, Phe122, Glu123, Leu126, and Ser127 throughout the entire simulation. Upon phosphorylation, these interactions are reduced. Additionally, Thr92 forms new, transient contacts with Pro124, Ser125, and Val128 when phosphorylated.

| | **Unph. CD36** | | **Ph. CD36** | |
|---|---|---|---|---|
| **Thr92** | Ser127 | 1.0 | Ser127 | 0.360 |
| | Leu126 | 1.0 | Leu126 | 0.246 |
| | Glu123 | 1.0 | Glu123 | 0.563 |
| | Phe122 | 1.0 | Phe122 | 0.897 |
| | Arg63 | 1.0 | Arg63 | 0.999 |
| | Tyr62 | 0.999 | Tyr62 | 0.934 |
| | Gln64 | 0.997 | Gln64 | 0.939 |
| | Ile121 | 0.561 | Ile121 | 0.361 |
| | | | Ser125 | 0.284 |
| | | | Pro124 | 0.104 |
| | | | Val128 | 0.066 |
| **Ser237** | Asp244 | 0.886 | Asp244 | 0.999 |
| | Lys233 | 0.700 | Lys233 | 0.996 |

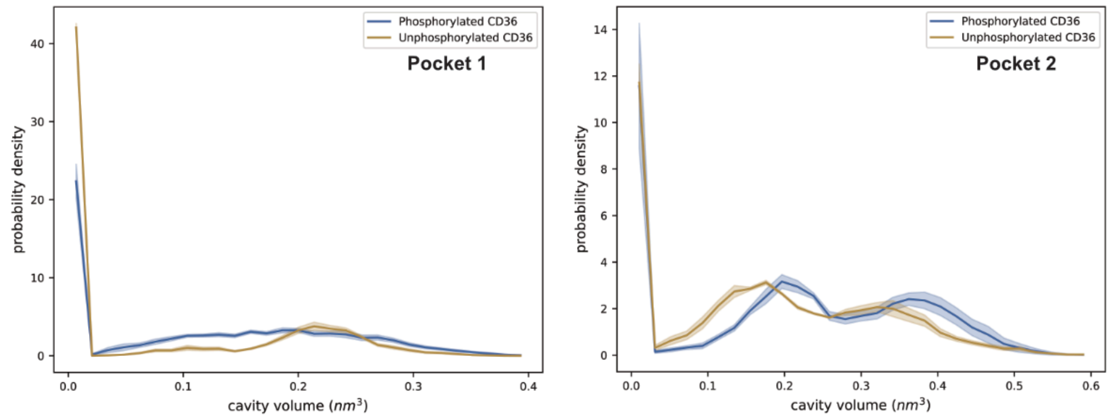


**Figure S9.** Distribution of the volume of pockets 1 and 2 averaged over five, 3 µs-long trajectories. Shading indicates uncertainty (standard error of the mean) over the five replicates.

Table S3. CD36 residues forming pocket 3 in the phosphorylated state.

| Residue type | Residue number |
|---|---|
| Tyr | 62 |
| Gln | 64 |
| p-Thr | 92 |
| Tyr | 93 |
| Arg | 94 |
| Ile | 121 |
| Phe | 122 |
| Glu | 123 |
| Pro | 124 |
| Ser | 125 |
| Leu | 126 |
| Thr | 419 |

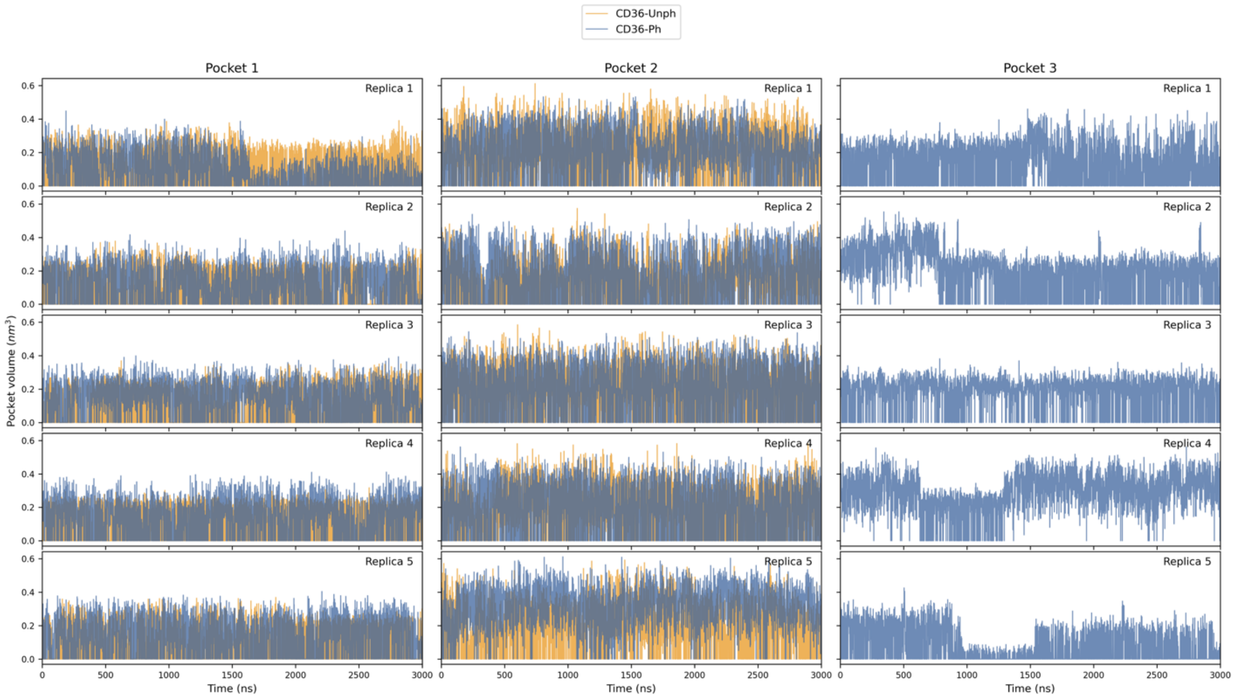


**Figure S10.** Time series of pocket 1, 2, and 3 volumes throughout the 3 μs simulation period for five independent simulations of unphosphorylated and phosphorylated CD36. Pocket 3 is observed only in phosphorylated CD36.

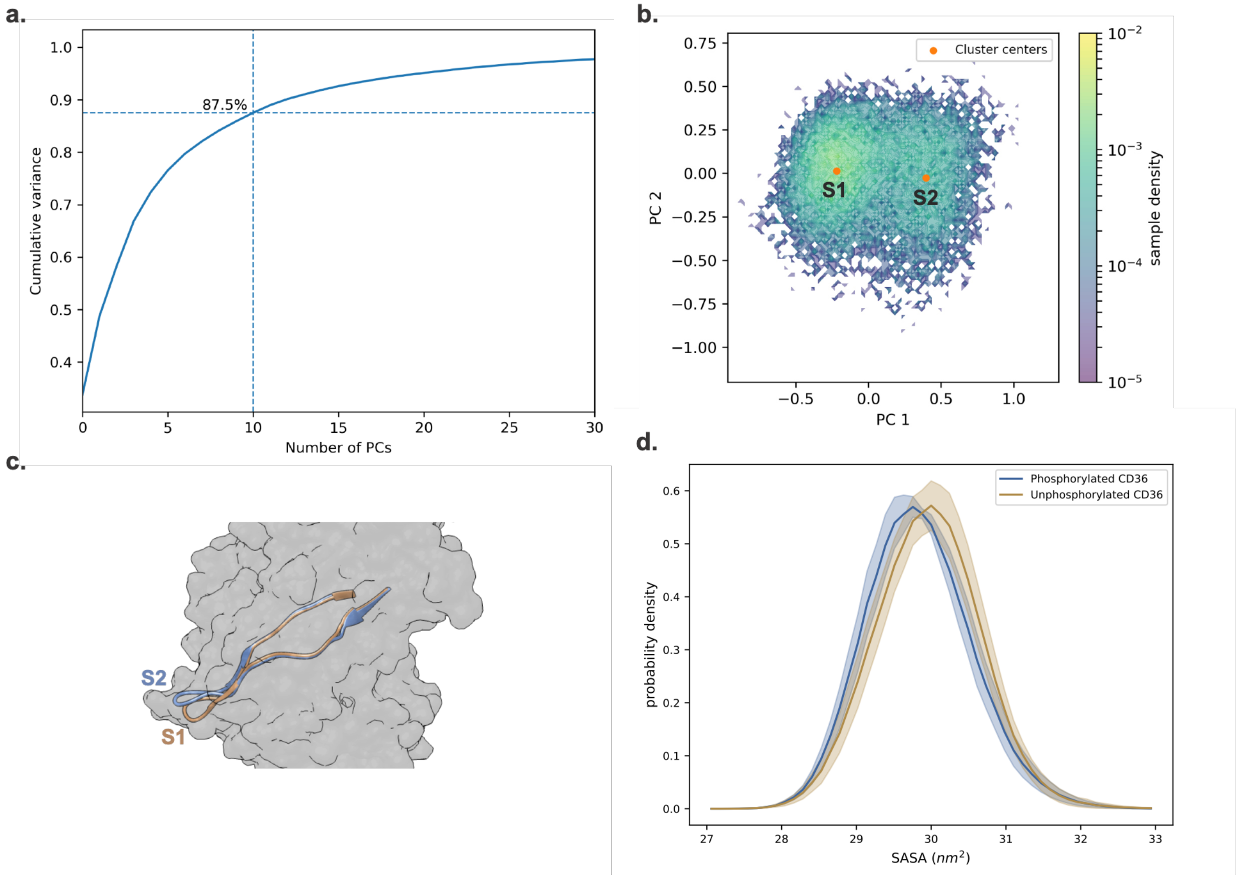


**Figure S11.** PCA and clustering results for the phosphorylated CD36 TSP binding site (CLESH domain: residues 93-120). (a) Proportion of variance explained by the principal components (PCs); the first ten PCs account for 87.5% of the total variance. (b) Density plot of the data projected onto the first two PCs, with clusters centers shown in orange dots. (c) Representative structures from each cluster overlaid for comparison. (d) Solvent accessible surface area (SASA) of CLESH domain in CD36 structure: phosphorylated vs. unphosphorylated forms.

**Table S4.** The hydropathicity of the peptide ligands derived from the TSR2 domain of TSP-1. Residues are color-coded based on their charge and hydropathic character: blue for positively charged, red for negatively charged, green for hydrophilic, and khaki for hydrophobic residues. The hydropathic character is assigned using the Kyte and Doolittle hydrophobicity scale (kdHydrophobicity) [1], where positive values indicate hydrophobic residues and negative values indicate hydrophilic residues.

| Ligand 1 (TSR2 residues 418-443) | | Ligand 2 (TSR2 residues 438-464) | |
|---|---|---|---|
| **Residue** | **kdHydrophobicity** | **Residue** | **kdHydrophobicity** |
| Gly 418 | -0.4 | Ile 438 | 4.5 |
| Gly 419 | -0.4 | Thr 439 | -0.7 |
| Trp 420 | -0.9 | Arg 440 | -4.5 |
| Ser 421 | -0.8 | Ile 441 | 4.5 |
| His 422 | -3.2 | Arg 442 | -4.5 |
| Trp 423 | -0.9 | Leu 443 | 3.8 |
| Ser 424 | -0.8 | Cys 444 | 2.5 |
| Pro 425 | -1.6 | Asn 445 | -3.5 |
| Trp 426 | -0.9 | Ser 446 | -0.8 |
| Ser 427 | -0.8 | Pro 447 | -1.6 |
| Ser 428 | -0.8 | Ser 448 | -0.8 |
| Cys 429 | 2.5 | Pro 449 | -1.6 |
| Ser 430 | -0.8 | Gln 450 | -3.5 |
| Val 431 | 4.2 | Met 451 | 1.9 |
| Thr 432 | -0.7 | Asn 452 | -3.5 |
| Cys 433 | 2.5 | Gly 453 | -0.4 |
| Gly 434 | -0.4 | Lys 454 | -3.9 |
| Asp 435 | -3.5 | Pro 455 | -1.6 |
| Gly 436 | -0.4 | Cys 456 | 2.5 |
| Val 437 | 4.2 | Glu 457 | -3.5 |
| Ile 438 | 4.5 | Gly 458 | -0.4 |
| Thr 439 | -0.7 | Glu 459 | -3.5 |
| Arg 440 | -4.5 | Ala 460 | 1.8 |
| Ile 441 | 4.5 | Arg 461 | -4.5 |
| Arg 442 | -4.5 | Glu 462 | -3.5 |
| Leu 443 | 3.8 | Thr 463 | -0.7 |
| - | - | Lys 464 | -3.9 |
| **Total score** | **-0.8** | | **-29.4** |

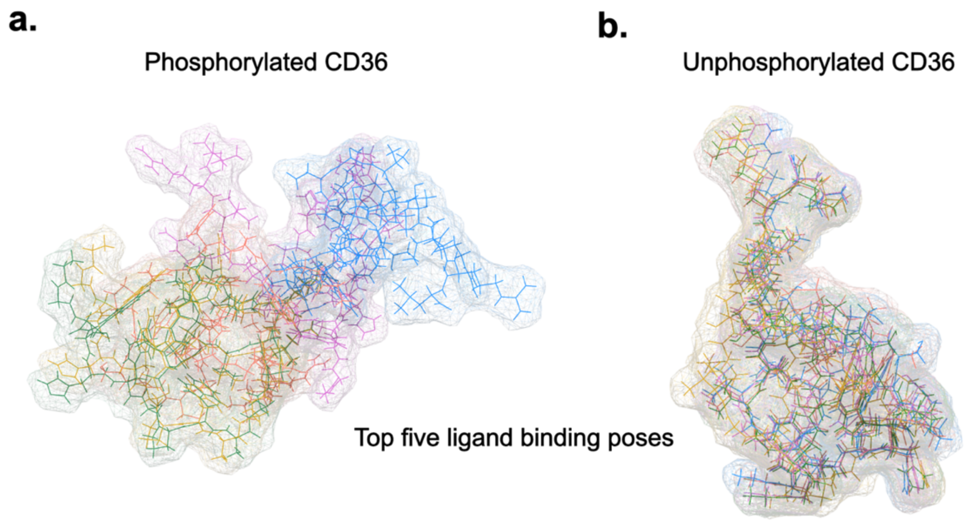


**Figure S12.** Top five ligand binding poses of the peptide ligand 1 derived from TSP-1 type 1 repeat 2 (TSR2) (residues 418-443) to CD36 receptor is shown for phosphorylated and unphosphorylated CD36.

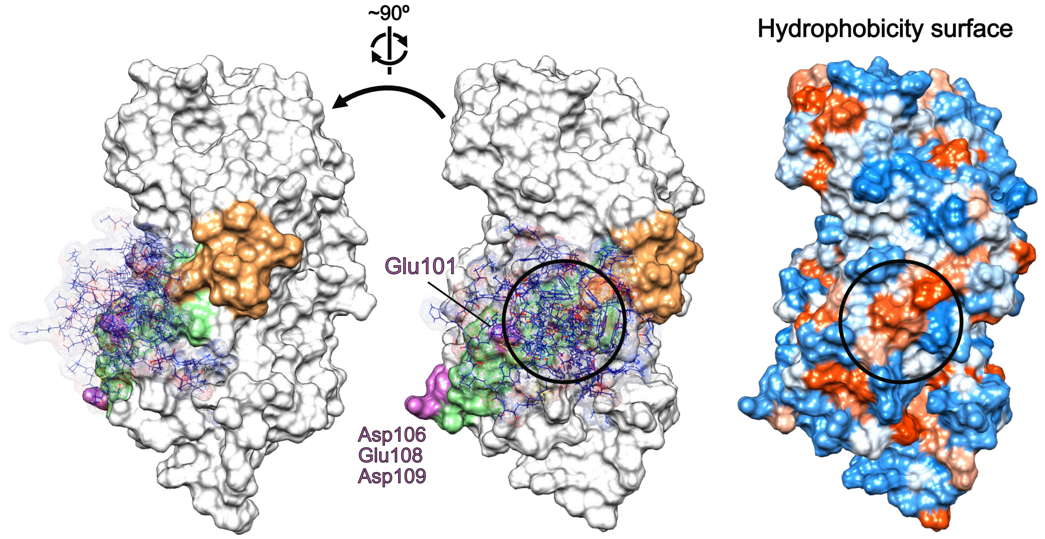


**Figure S13.** Less favored ensemble docking result of the peptide ligand 1 derived from TSR2 (residues 418-443) to unphosphorylated CD36 structure. CD36 structure and top five ligand binding poses are shown in solid surface and wire representation, respectively. Reg2 and the TSP binding site are highlighted in orange and green, respectively. Key residues of the CLESH domain (Glu101, Asp106, Glu108, and Asp109) are highlighted in purple. The hydrophobic surface of unphosphorylated CD36 is depicted on the right, with color intensity ranging from red (indicating the most hydrophobic regions) to blue (indicating the most hydrophilic regions). Black circle indicates the approximate binding area of the top 5 ligand poses.

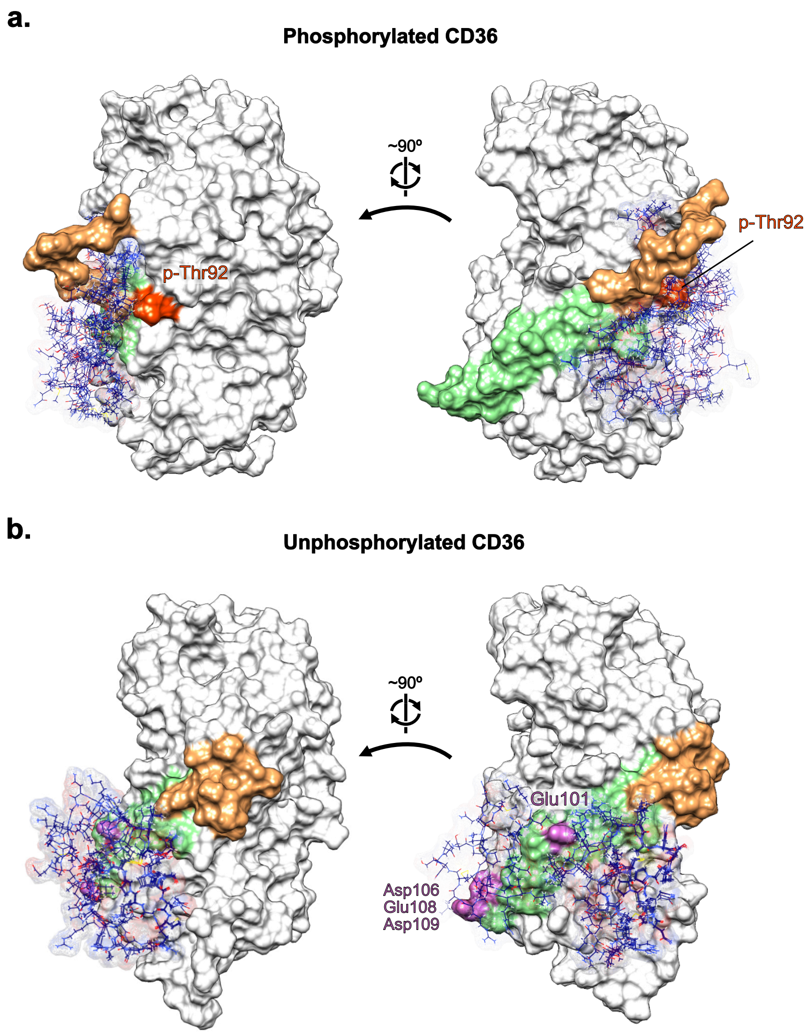


**Figure S14.** Ensemble docking results of the peptide ligand 2 derived from TSR2 (residues 438-464) to CD36 receptor is shown for (a) phosphorylated and (b) unphosphorylated CD36. CD36 structure and top five ligand binding poses are shown in solid surface and wire representation, respectively. Reg2 and the TSP binding site are highlighted in orange and green, respectively. Key residues of the CLESH domain (Glu101, Asp106, Glu108, and Asp109) are highlighted in purple within the unphosphorylated CD36. The phosphorylated Thr92 is depicted in red.

**Table S5.** CD36 residues that are in contact with (within 5Å) the TSR2 ligands for the top five docking poses. Residues are color-coded based on their charge and hydropathic character: blue for positively charged residues, red for negatively charged residues, green for hydrophilic residues, and khaki for hydrophobic residues. The hydropathic character is assigned using the Kyte and Doolittle hydrophobicity scale (kdHydrophobicity) [1], where positive values indicate hydrophobic residues and negative values indicate hydrophilic residues. The orange highlighted part indicates residues from the region spanning residues 92 to 131, which includes Thr92, the CLESH domain and Reg2.

| Unphosphorylated CD36 | | | | | |
|---|---|---|---|---|---|
| Ligand 1 (TSR2 residues 418-443) | | | | Ligand 2 (TSR2 residues 438-464) | |
| In 8 out of 10 docking models | | In 2 out of 10 docking models | | | |
| Residue | kdHydrophobicity | Residue | kdHydrophobicity | Residue | kdHydrophobicity |
| | | | | LYS 36 | -3.9 |
| LYS 40 | -3.9 | | | | |
| | | GLN 41 | -3.5 | | |
| GLU 45 | -3.5 | | | GLU 45 | -3.5 |
| GLU 46 | -3.5 | | | GLU 46 | -3.5 |
| GLY 47 | -0.4 | | | | |
| THR 48 | -0.7 | | | THR 48 | -0.7 |
| | | ILE 49 | 4.5 | ILE 49 | 4.5 |
| | | ALA 50 | 1.8 | ALA 50 | 1.8 |
| PHE 51 | 2.8 | PHE 51 | 2.8 | PHE 51 | 2.8 |
| LYS 52 | -3.9 | LYS 52 | -3.9 | LYS 52 | -3.9 |
| | | ASN 53 | -3.5 | ASN 53 | -3.5 |
| | | TRP 54 | -0.9 | | |
| | | VAL 55 | 4.2 | VAL 55 | 4.2 |
| | | LYS 56 | -3.9 | LYS 56 | -3.9 |
| | | THR 57 | -0.7 | THR 57 | -0.7 |
| | | GLY 58 | -0.4 | GLY 58 | -0.4 |
| | | THR 59 | -0.7 | | |
| | | GLU 60 | -3.5 | | |
| | | VAL 61 | 4.2 | | |
| | | ARG 94 | -4.5 | ARG 94 | -4.5 |
| | | ARG 96 | -4.5 | | |
| | | PHE 97 | 2.8 | PHE 97 | 2.8 |
| | | LEU 98 | 3.8 | LEU 98 | 3.8 |
| | | ALA 99 | 1.8 | ALA 99 | 1.8 |
| | | LYS 100 | -3.9 | LYS 100 | -3.9 |
| GLU 101 | -3.5 | GLU 101 | -3.5 | GLU 101 | -3.5 |
| ASN 102 | -3.5 | | | ASN 102 | -3.5 |
| VAL 103 | 4.2 | | | VAL 103 | 4.2 |
| THR 104 | -0.7 | | | THR 104 | -0.7 |
| GLN 105 | -3.5 | | | GLN 105 | -3.5 |
| ASP 106 | -3.5 | | | ASP 106 | -3.5 |
| ALA 107 | 1.8 | | | ALA 107 | 1.8 |
| GLU 108 | -3.5 | | | GLU 108 | -3.5 |
| ASP 109 | -3.5 | | | | |
| THR 111 | -0.7 | | | | |
| | | | | VAL 112 | 4.2 |
| SER 113 | -0.8 | | | SER 113 | -0.8 |
| PHE 114 | 2.8 | | | | |
| LEU 115 | 3.8 | | | LEU 115 | 3.8 |
| | | PRO 117 | -1.6 | PRO 117 | -1.6 |
| | | ASN 118 | -3.5 | ASN 118 | -3.5 |
| | | GLY 119 | -0.4 | GLY 119 | -0.4 |
| | | ALA 120 | 1.8 | ALA 120 | 1.8 |
| | | ILE 121 | 4.5 | ILE 121 | 4.5 |
| | | ARG 176 | -4.5 | | |

| | | | | | |
|---|---|---|---|---|---|
| | | ASN 205 | -3.5 | | |
| | | ASN 206 | -3.5 | | |
| | | THR 207 | -0.7 | THR 207 | -0.7 |
| | | ALA 208 | 1.8 | ALA 208 | 1.8 |
| | | | | GLY 210 | -0.4 |
| VAL 211 | 4.2 | | | VAL 211 | 4.2 |
| TYR 212 | -1.3 | | | | |
| LYS 213 | -3.9 | | | LYS 213 | -3.9 |
| ASN 216 | -3.5 | | | | |
| LYS 218 | -3.9 | | | | |
| LYS 231 | -3.9 | | | LYS 231 | -3.9 |
| GLY 232 | -0.4 | | | | |
| LYS 233 | -3.9 | | | | |
| | | ALA 252 | 1.8 | | |
| | | THR 377 | -0.7 | | |
| | | PHE 379 | 2.8 | | |
| | | LEU 381 | 3.8 | | |
| | | PHE 383 | 2.8 | | |
| | | ILE 422 | 4.5 | | |
| | | GLY 423 | -0.4 | | |
| | | | | GLU 425 | -3.5 |
| | | LYS 426 | -3.9 | LYS 426 | -3.9 |
| | | ALA 427 | 1.8 | | |
| | | MET 429 | 1.9 | MET 429 | 1.9 |
| | | PHE 430 | 2.8 | PHE 430 | 2.8 |
| | | ARG 431 | -4.5 | | |
| | | | | SER 432 | -0.8 |
| | | GLN 433 | -3.5 | GLN 433 | -3.5 |
| **Total score** | **-40.3** | **Total score** | **-11.9** | **Total score** | **-24.8** |
| **Phosphorylated CD36** | | | | | |
| **Ligand 1 (TSR2 residues 418-443)** | | | | **Ligand 2 (TSR2 residues 438-464)** | |
| **Residue** | **kdHydrophobicity** | | | **Residue** | **kdHydrophobicity** |
| ILE 49 | 4.5 | | | | |
| | | | | LYS 52 | -3.9 |
| ASN 53 | -3.5 | | | ASN 53 | -3.5 |
| VAL 55 | 4.2 | | | | |
| LYS 56 | -3.9 | | | LYS 56 | -3.9 |
| | | | | THR 57 | -0.7 |
| GLY 58 | -0.4 | | | GLY 58 | -0.4 |
| THR 59 | -0.7 | | | THR 59 | -0.7 |
| GLU 60 | -3.5 | | | GLU 60 | -3.5 |
| TYR 62 | -1.3 | | | TYR 62 | -1.3 |
| | | | | ILE 67 | 4.5 |
| PRO 90 | -1.6 | | | | |
| TYR 91 | -1.3 | | | TYR 91 | -1.3 |
| p-THR 92 | Hydrophilic | | | p-THR 92 | Hydrophilic |
| TYR 93 | -1.3 | | | TYR 93 | -1.3 |
| ARG 94 | -4.5 | | | ARG 94 | -4.5 |
| PHE 97 | 2.8 | | | PHE 97 | 2.8 |
| LEU 98 | 3.8 | | | LEU 98 | 3.8 |
| ALA 99 | 1.8 | | | ALA 99 | 1.8 |
| LYS 100 | -3.9 | | | | |
| GLU 101 | -3.5 | | | | |
| PRO 117 | -1.6 | | | | |
| ASN 118 | -3.5 | | | | |
| GLY 119 | -0.4 | | | | |
| ALA 120 | 1.8 | | | ALA 120 | 1.8 |
| ILE 121 | 4.5 | | | ILE 121 | 4.5 |

| | | | | | | | |
|---|---|---|---|---|---|---|---|
| PHE | 122 | 2.8 | | | PHE | 122 | 2.8 |
| GLU | 123 | -3.5 | | | GLU | 123 | -3.5 |
| PRO | 124 | -1.6 | | | PRO | 124 | -1.6 |
| SER | 125 | -0.8 | | | SER | 125 | -0.8 |
| LEU | 126 | 3.8 | | | LEU | 126 | 3.8 |
| SER | 127 | -0.8 | | | SER | 127 | -0.8 |
| VAL | 128 | 4.2 | | | VAL | 128 | 4.2 |
| GLY | 129 | -0.4 | | | GLY | 129 | -0.4 |
| THR | 130 | -0.7 | | | THR | 130 | -0.7 |
| GLU | 131 | -3.5 | | | GLU | 131 | -3.5 |
| ALA | 132 | 1.8 | | | ALA | 132 | 1.8 |
| ASP | 133 | -3.5 | | | ASP | 133 | -3.5 |
| | | | | | ASN | 134 | -3.5 |
| | | | | | PHE | 135 | 2.8 |
| | | | | | THR | 174 | -0.7 |
| LEU | 175 | 3.8 | | | LEU | 175 | 3.8 |
| ARG | 176 | -4.5 | | | ARG | 176 | -4.5 |
| GLU | 177 | -3.5 | | | GLU | 177 | -3.5 |
| TRP | 180 | -0.9 | | | | | |
| GLY | 181 | -0.4 | | | | | |
| ASN | 205 | -3.5 | | | | | |
| ASN | 206 | -3.5 | | | | | |
| THR | 421 | -0.7 | | | | | |
| ILE | 422 | 3.8 | | | | | |
| GLY | 423 | -0.4 | | | GLY | 423 | -0.4 |
| ASP | 424 | -3.5 | | | ASP | 424 | -3.5 |
| GLU | 425 | -3.5 | | | GLU | 425 | -3.5 |
| LYS | 426 | -3.9 | | | LYS | 426 | -3.9 |
| ALA | 427 | 1.8 | | | | | |
| **Total score** | | **-33*** | | | **Total score** | | **-24.9*** |

* Without considering the hydrophilicity of p-Thr92.

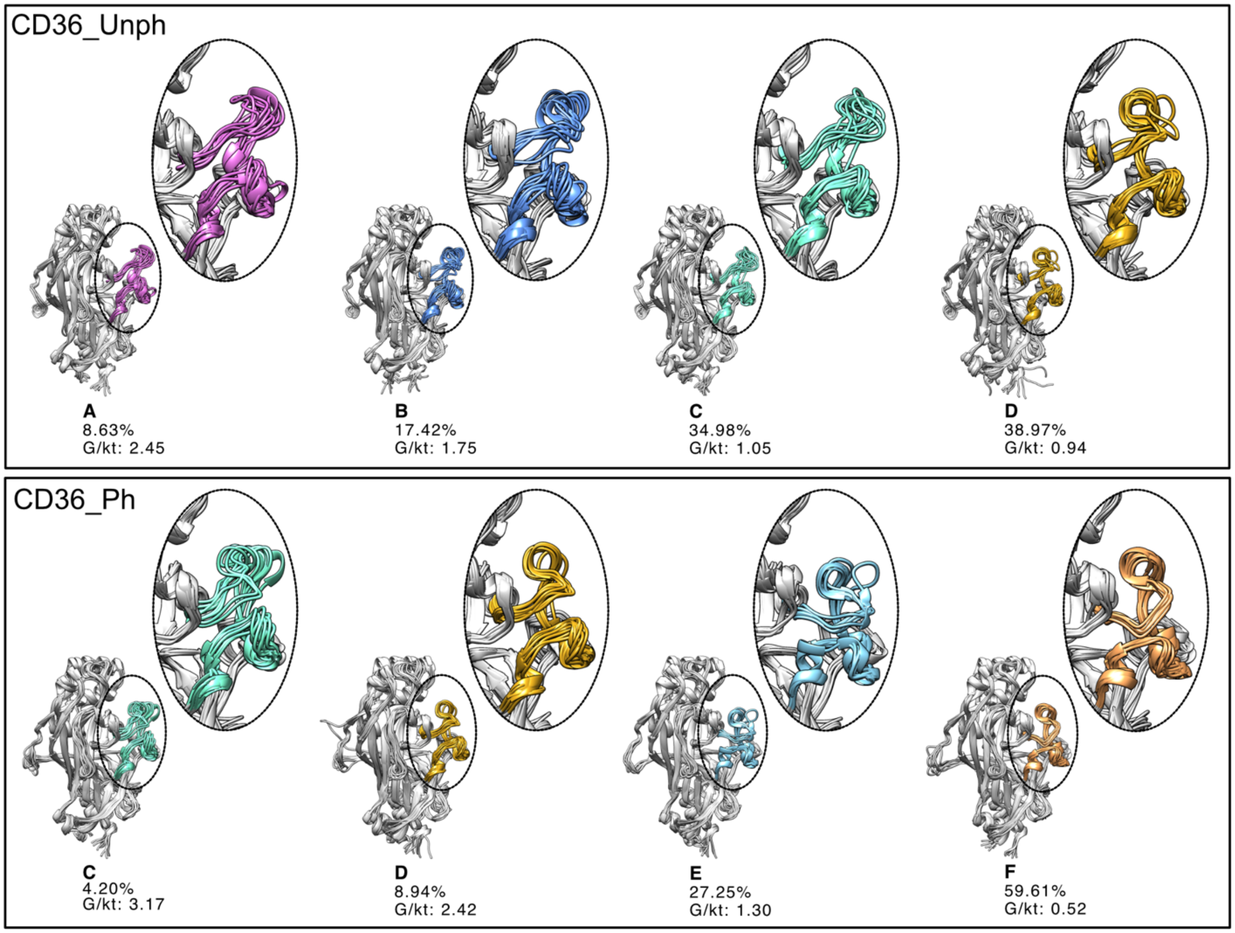


**Figure S15.** Ten randomly selected representative conformations of Reg1 (residues 296-331) for each metastable state from a 15 μs ensemble for unphosphorylated (top) and phosphorylated CD36 (bottom) systems.

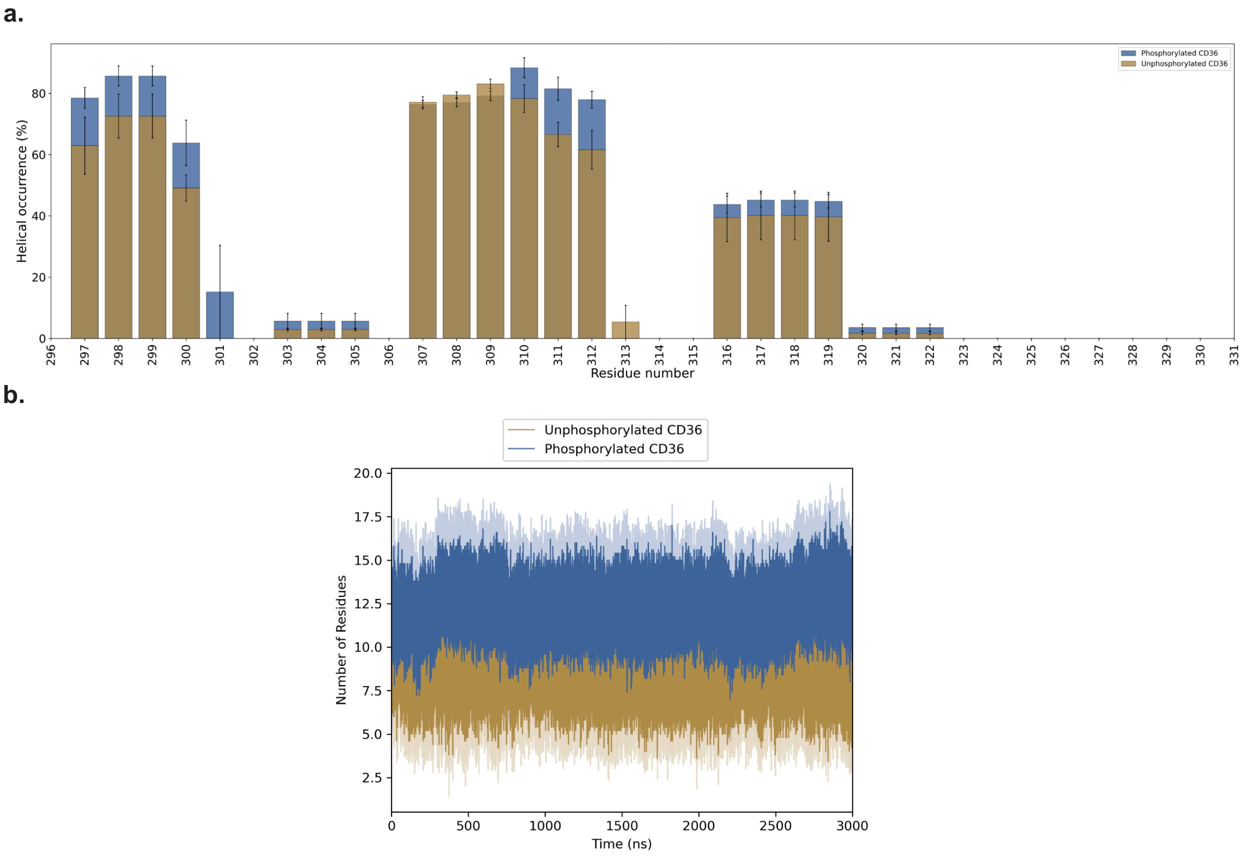


**Figure S16.** Helical structure occurrence in Reg1 (residues 296-331) for phosphorylated and unphosphorylated CD36. (a) Percentage of helical occurrence for each residue in Reg1, averaged across five independent, 3 µs-long trajectories. (b) Number of residues forming a helical structure throughout the 3 µs simulation period. Error bars in (a) and the shading in (b) indicate the standard error of the mean over five independent simulations.

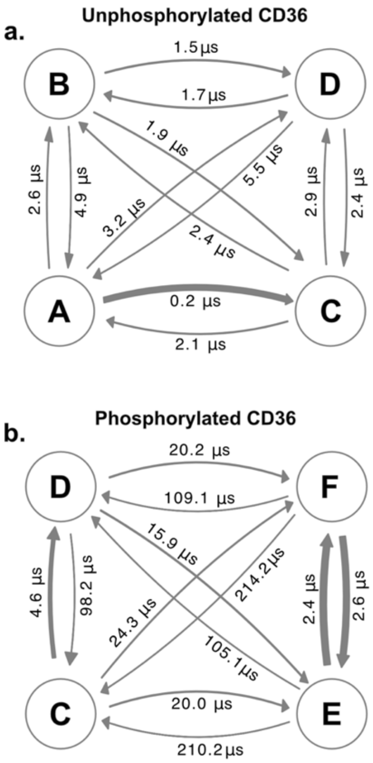


**Figure S17.** Kinetic model showing the mean first passage time (MFPT) across states for Reg1 in (a) unphosphorylated and (b) phosphorylated CD36.

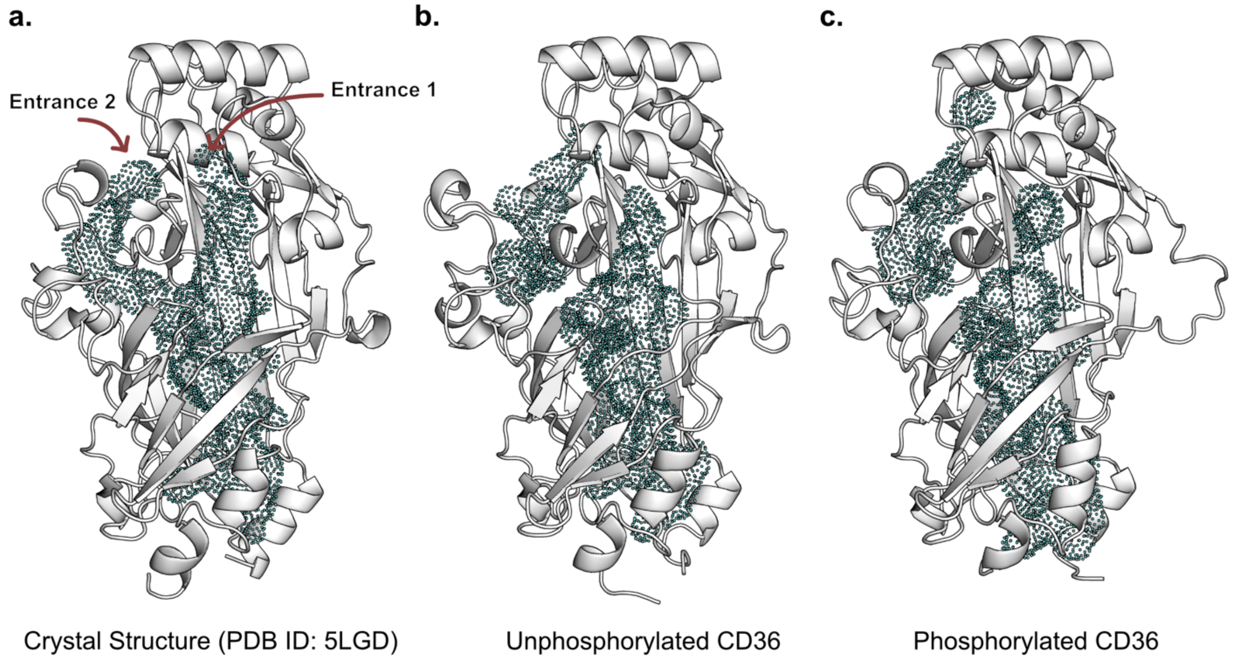


**Figure S18.** Hydrophobic cavity inside CD36 structure in the (a) crystal structure, (b) unphosphorylated and (c) phosphorylated forms. Analyzing eight randomly chosen structures of both unphosphorylated and phosphorylated CD36, each representing a metastable state from the Reg1 MSMs, shows that Entrance 2 and the rest of the internal cavity are disconnected. This suggests that the dynamics of the protein can affect the size and shape of the internal cavity irrespective of the phosphorylation state of the protein. Computed Atlas of Surface Topography of proteins (CASTp) server [2] was used to identify the internal hydrophobic cavity of the CD36 structure, with a default value of 1.4 Å for the probe radius. PyMOL (version 2.5.0) was used to visualize the cavity.

**Table S6.** Force field parameters used for the phosphorylated residues, Thr92 and Ser237.

**Angles:**

| i | j | k | func | theta0 | ktheta | ub0 | kub |
|---|---|---|---|---|---|---|---|
| ON2 | P | ON2 | 5 | 104.300000 | 669.440000 | 0.00000000 | 0.00 |
| ON2 | P | ON4 | 5 | 108.000000 | 402.500800 | 0.00000000 | 0.00 |
| ON3 | P | ON4 | 5 | 108.230000 | 827.595200 | 0.00000000 | 0.00 |
| ON4 | P | ON4 | 5 | 104.000000 | 827.595200 | 0.00000000 | 0.00 |
| ON2 | P | ON3 | 5 | 111.600000 | 827.595200 | 0.00000000 | 0.00 |
| ON3 | P | ON3 | 5 | 120.000000 | 1004.160000 | 0.00000000 | 0.00 |

**Dihedrals:**

| i | j | k | l | func | phi0 | kphi | mult | Remark |
|---|---|---|---|---|---|---|---|---|
| ON3 | P | ON2 | CT2 | 9 | 0.0 | 0.4184 | 3 | Adopted from ON3 P ON2 CN8B |
| P | ON2 | CT2 | HA2 | 9 | 0.0 | 0.0 | 3 | Adopted from P ON2 CN8B HN8 |

**Following parameters were adopted from CHARMM36:**

**Bonds:**

| i | j | func | b0 | kb | Remark |
|---|---|---|---|---|---|
| CT1 | ON2 | 1 | 0.14330000 | 259408.00 | Adopted from CN7 ON2<br>CT1 12.01100 ; aliphatic sp3 C for CH<br>CN7 12.011000 ; Nucleic acid carbon (equivalent to protein CT1) ### For DNA |
| CT2 | ON2 | 1 | 0.14400000 | 267776.00 | Adopted from CN8B ON2<br>CN8 12.011000 ; Nucleic acid carbon (equivalent to protein CT2) ### For DNA<br>CN8B 12.011000 ; Nucleic acid carbon (equivalent to protein CT2) ### For DNA |

**Angles:**

| i | j | k | func | theta0 | ktheta | ub0 | kub | Remark |
|---|---|---|---|---|---|---|---|---|
| CT1 | CT1 | ON2 | 5 | 109.700000 | 962.320000 | 0.00000000 | 0.00 | Adopted from CN7 CN7 ON2 |
| HA1 | CT1 | ON2 | 5 | 109.500000 | 502.080000 | 0.00000000 | 0.00 | Adopted from HN7 CN7 ON2 |
| ON2 | CT1 | CT3 | 5 | 109.700000 | 962.320000 | 0.00000000 | 0.00 | Adopted from CN8 CN7 ON2 |
| CT1 | ON2 | P | 5 | 120.000000 | 167.360000 | 0.23300000 | 29288.00 | Adopted from CN7 ON2 P |
| CT1 | CT2 | ON2 | 5 | 108.400000 | 585.760000 | 0.00000000 | 0.00 | Adopted from CN7 CN8B ON2 |
| HA2 | CT2 | ON2 | 5 | 109.500000 | 502.080000 | 0.00000000 | 0.00 | Adopted from HN8 CN8B ON2 |
| CT2 | ON2 | P | 5 | 120.000000 | 167.360000 | 0.23300000 | 29288.00 | Adopted from CN8B ON2 P |

**Dihedrals:**

| I | J | k | l | func | phi0 | kphi | mult | Remark |
|---|---|---|---|---|---|---|---|---|
| CT1 | CT1 | ON2 | P | 9 | 0.000000<br>0.000000<br>180.000000<br>0.000000<br>180.000000 | 2.510400<br>0.836800<br>0.000000<br>1.673600<br>7.949600 | 5<br>4<br>3<br>2<br>1 | Adopted from CN7 CN7 ON2 P |
| HA1 | CT1 | ON2 | P | 9 | 0.000000 | 0.000000 | 3 | Adopted from P ON2 CN7 HN7 |
| CT3 | CT1 | ON2 | P | 9 | 180.000000 | 10.460000 | 1 | Adopted from the following:<br>CN7B CN7 ON2 P<br>CN7B CN7B ON2 P<br>CN7 CN7B ON2 P<br>CN8 CN7B ON2 P<br>CN7C CN7 ON2 P |
| CT1 | ON2 | P | ON3 | 9 | 0.000000 | 0.418400 | 3 | Adopted from ON3 P ON2 CN7 |
| CT1 | CT2 | ON2 | P | 9 | 120.000000 | 0.836800 | 1 | Adopted from CN7 CN8B ON2 P |
| CT2 | ON2 | P | ON3 | 9 | 0.000000 | 0.418400 | 3 | Adopted from ON3 P ON2 CN8B |
| P | ON2 | CT2 | HA2 | 9 | 0.000000 | 0.000000 | 3 | Adopted from P ON2 CN8B HN8 |

**Table S7.** Feature data refinement steps for MSM construction

| **Steps** | **Number of resulting features from each step** | **Lag time** | **Number of tICs** |
|---|---|---|---|
| Calculate all possible $C_\alpha$-$C_\alpha$ distances | 79,003 | - | - |
| Remove distances < 3 Å and > 10 Å | 5,154 | - | - |
| Remove data with variance < 0.005 | 1,441 | - | - |
| Remove distances involving terminal residues | 1,372 | - | - |
| Start iterative TICA-based data removal: | 1,372 | 10 ns | 678 |
| Remove data with correlation < 0.4 | 695 | 10 ns | 307 |
| Remove data with correlation < 0.6 | 377 | 10 ns | 137 |
| Remove data with correlation < 0.7 | 212 | 10 ns | 66 |
| Remove data with correlation < 0.8 | 110 | 10 ns | 36 |
| Remove data with correlation < 0.85 | 61 | 10 ns | 23 |
| Remove data with correlation < 0.9 | 26 | - | - |
| Add features identified in unphosphorylated CD36 system | 38 | - | - |

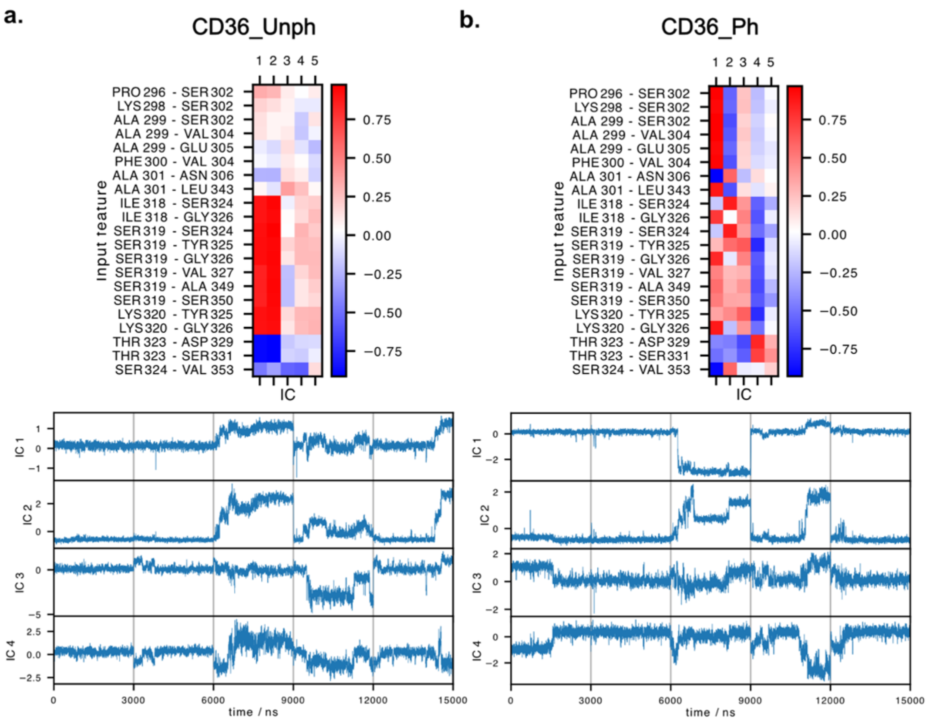


**Figure S19.** Projection of the Reg1 results for the (a) unphosphorylated and (b) phosphorylated CD36 system. The top figures show the correlation between the 21 selected features and the first five ICs (lag=10). The bottom figures show all the trajectories projected on the first 4 ICs. Each trajectory is divided by a gray line.

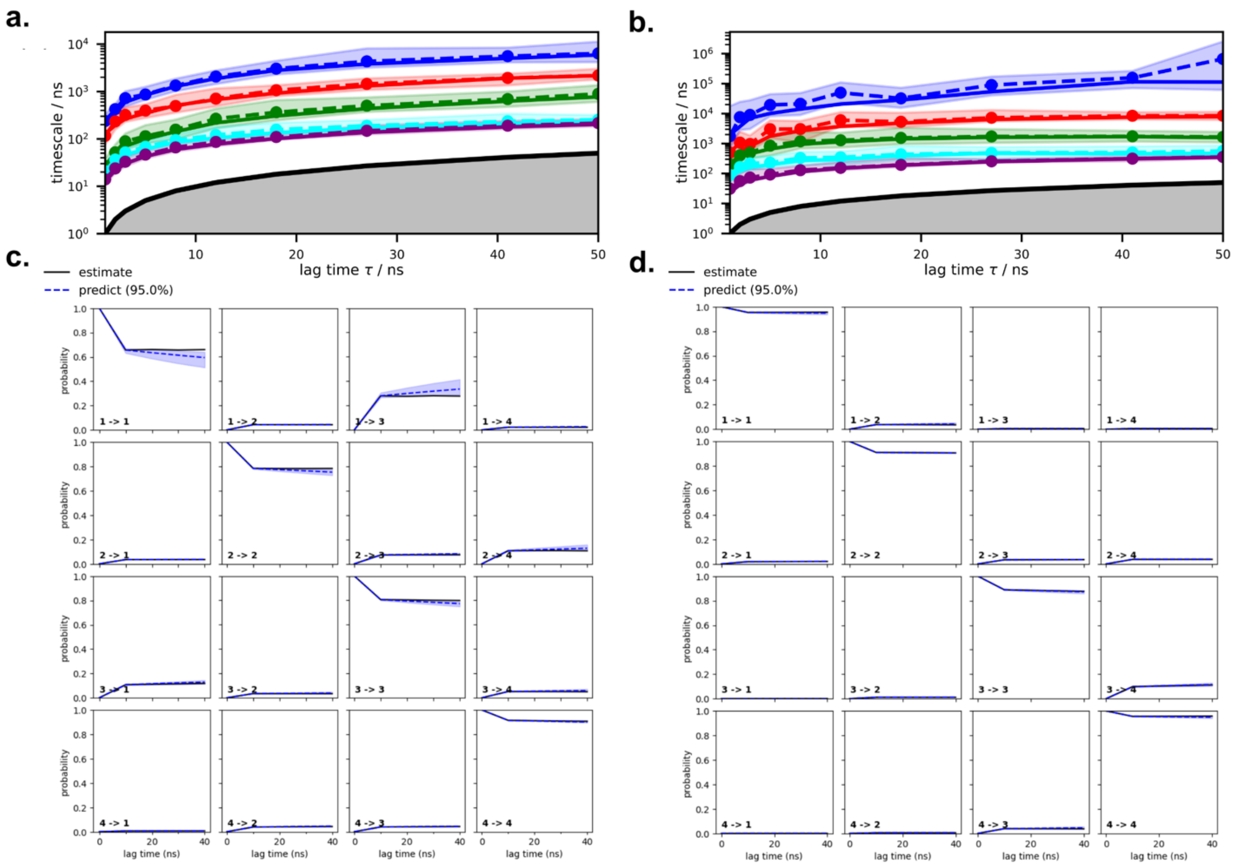


**Figure S20.** Lag-time determination and validation of MSMs for Reg1. Independent time scales for (a) unphosphorylated and (b) phosphorylated CD36 systems. The selected lag time was 10 ns for MSM estimation. Chapman−Kolmogorov test result for (c) unphosphorylated and (d) phosphorylated CD36 systems.

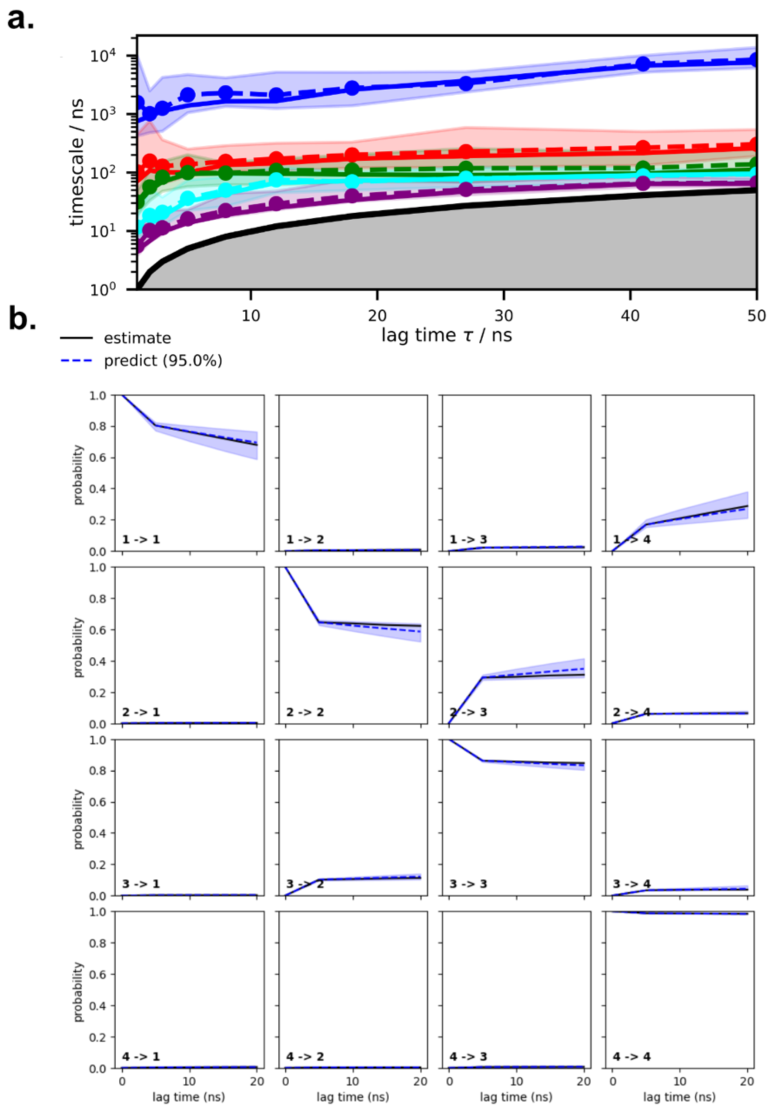


**Figure S21.** Lag-time determination and validation of MSM for Reg2. (a) Independent time scales for phosphorylated CD36 system. The selected lag time was 5 ns for MSM estimation. (b) Chapman−Kolmogorov test result for phosphorylated CD36 system.

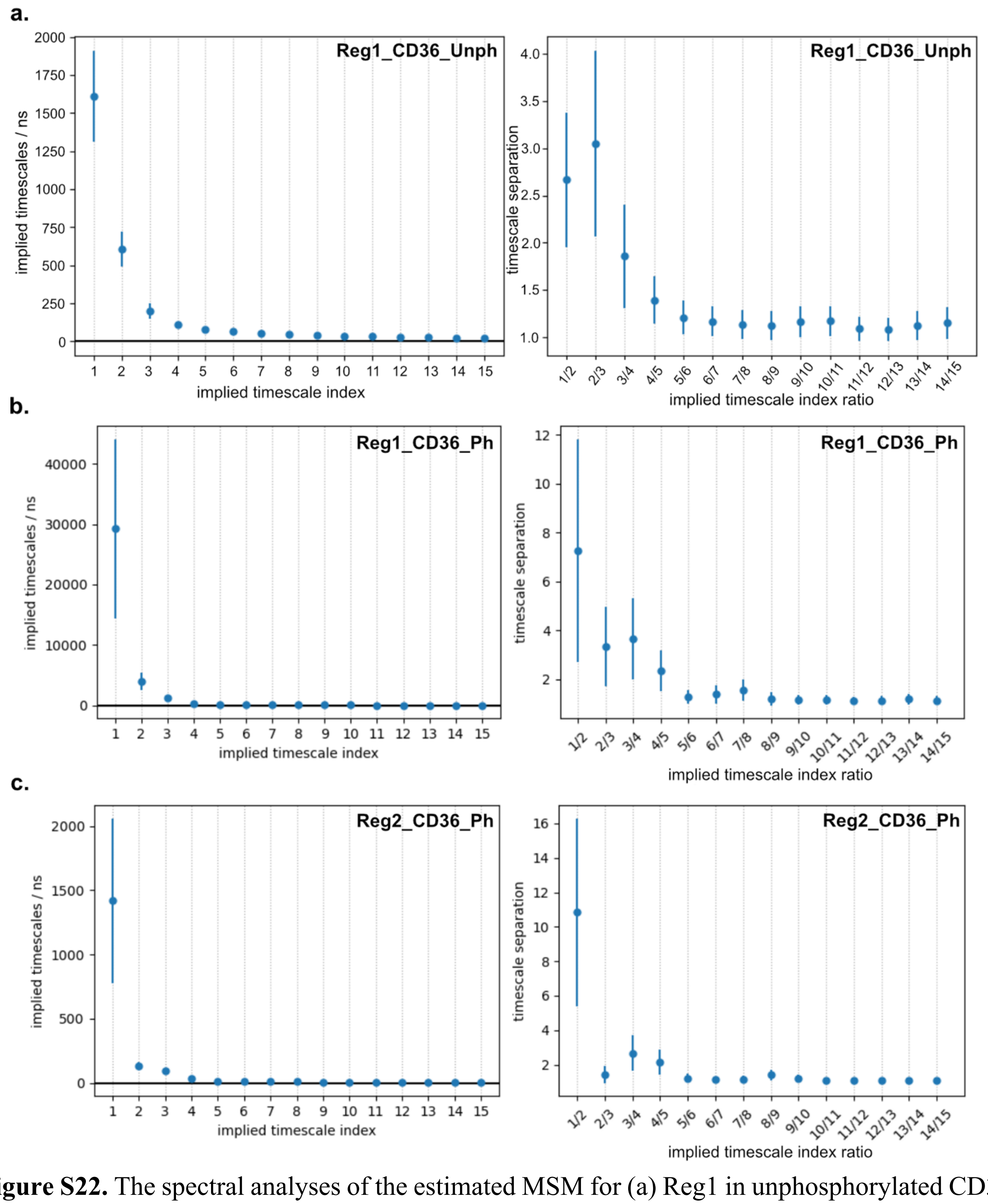


**Figure S22.** The spectral analyses of the estimated MSM for (a) Reg1 in unphosphorylated CD36, (b) Reg1 in phosphorylated CD36 and (c) Reg2 in phosphorylated CD36. The timescale separation between the 3rd and 4th process confirms that 4 metastable states is a suitable choice for coarse graining the microstates.

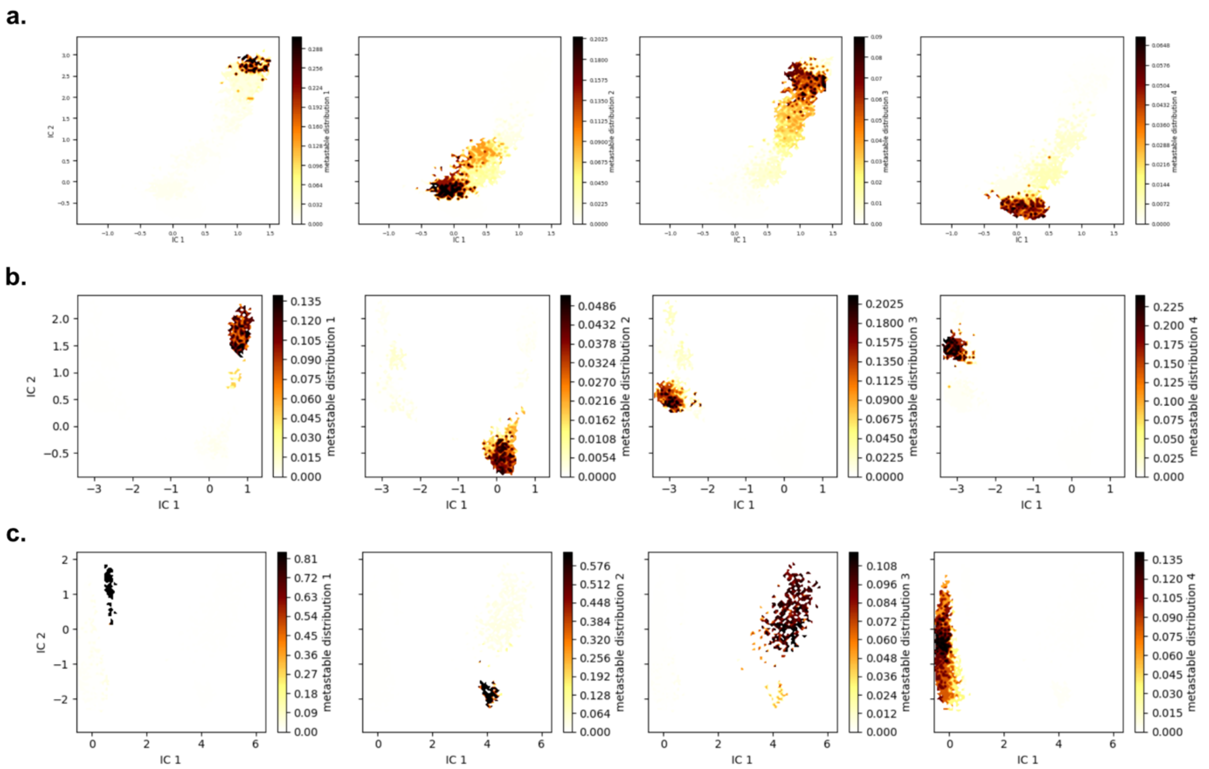

**Figure S23.** Membership graphs showing the probability of each microstate to belong to a given macrostate for (a) Reg1 in unphosphorylated CD36, (b) Reg1 and (c) Reg2 in phosphorylated CD36.
